# Validation of tissue-specific RNAi systems in *C. elegans* reveals a converging role for polyubiquitin UBQ-1/UBC in vitellogenin metabolism and lifespan

**DOI:** 10.64898/2026.02.21.707171

**Authors:** Noeli Soares Melo da Silva, Cristina Bolonyi, Adia Ouellette, Laura Harrison, Sung Youn Kim, Sandrine Daigle, Savannah T. Doucet, Louis R. Lapierre

## Abstract

Numerous studies in *C. elegans* have taken advantage of gene silencing RNAi libraries and tissue-specific RNAi systems to temporally and spatially understand gene function. Tissue-specific RNAi strains were created via tissue-specific functional expression of a key gene in the RNAi system in a presumably RNAi insensitive correspondingly mutated background. Here, we tested the level of RNAi insensitivity across different commonly used RNAi-defective mutants (*rde-1* and *sid-*1) using lifespan analyses. Remarkably, while we found similarly important lifespan shortening in wild-type animals by silencing the polyubiquitin gene *ubq-1* (*UBC*) at both 20°C and 25°C, we found wide and temperature-sensitive variations in RNAi sensitivity across different *rde-1* and *sid-1* strains from substantial sensitivity to complete insensitivity. This riveting finding warrants a re-evaluation of the tissue-specific interpretations in numerous tissue-specific RNAi studies in *C. elegans*. Using validated tissue-specific RNAi systems, we determined that proteasomal burden largely differs between the germline and the soma. Notably, gonadally accumulating vitellogenin proteins substantially contributed to the total polyubiquitinated proteins observed during aging. Interestingly, we found that UBQ-1 was essential to maintain the transcription of highly expressed genes, including vitellogenins, supporting a broad and key role for polyubiquitin beyond proteostasis. Altogether, we authenticated commonly used tissue-specific RNAi systems and uncovered a new approach to improve RNAi insensitivity in RNAi-defective strains. With validated tissue-specific RNAi strains, we spatially analyzed proteasomal function and age-related polyubiquitinated protein accumulation and revealed a converging regulatory link between UBQ-1, vitellogenin metabolism and lifespan in *C. elegans*.

## INTRODUCTION

The advent of mRNA silencing with interfering RNA (RNAi) from bacterially-expressed double-stranded RNA empowered researchers to investigate specific and temporal control of gene expression in *C. elegans*, opening the door to studies of gene function across life stages [1]. This innovation also avoided ambiguous interpretations of adult-specific gene function when corresponding mutants have developmental issues. Genome-wide RNAi libraries from the Ahringer [2] and Vidal [3] laboratories were instrumental in providing broad and complementary coverage for gene silencing. Unbiased genome-wide RNAi screen, such as the ones performed to discover lifespan-modulating genes [4, 5], delivered an incredible wealth of information about genetic determinants of longevity and aging [6, 7].

Broad interest in understanding tissue-specific gene function in *C. elegans* eventually led to the development of strains that enable gene silencing in only one tissue. These strains were constructed in RNAi-defective mutants whereas functional reconstitution of the defective gene under a tissue-specific promoter would restore RNAi sensitivity in a tissue-specific manner. Three essential considerations needed to be met to allow tissue-specific RNAi strains to provide valid and rigorous interpretations of tissue-specific gene function. First, the RNAi-defective mutants needed to be entirely insensitive to RNAi as any remaining sensitivity will mislead interpretation. Mutations of the *rde-1* gene coding for a key protein in the RNA-induced silencing complex (RISC) complex [8] or mutations of the *sid-1* gene coding for a dsRNA receptor necessary for dsRNA import into tissues [9, 10] were generally selected to construct tissue-specific strains. Second, the tissue-specific promoter used to drive the expression of the functional gene had to be exclusively restricted to the tissue of interest [11]. Third, the 3’UTR regions in the expression plasmid needed to be compatible with the targeted tissue for reconstitution. This could be typically addressed by matching the promoter with the corresponding 3’UTR [12]. Ideally, such reconstitution would have restricted gene silencing in only a specific tissue and provided a tool to analyze tissue-specific gene function.

Clear warning signs emerged over the past decade about the rigor and reproducibility of tissue-specific RNAi strains developed in *C. elegans*, including not only the use of *rde-1* and *sid-1* mutant strains [13, 14] but also the use of mutants of the RNA-dependent polymerase *rrf-1* for germline-specific RNAi [15]. Regrettably, tissue-specific RNAi systems were initially designed on the unsubstantiated assumptions that RNAi-defectiveness equals completely insensitivity to RNAi and that certain promoters can drive the expression of function genes (such as RDE-1 or SID-1) in just one specific tissue [16]. Astonishingly, despite being used in hundreds of publications, systematic demonstration of the validity of different tissue-specific RNAi strains has been lacking.

One area of research where identifying the function of genes in a tissue-specific manner would meaningfully advance our understanding is the field of proteostasis and aging. For instance, complementary strategies including tissue-specific reconstitutions and recently-developed auxin-inducible degradation systems [17] have helped identify tissue-specific roles for the proteostasis-related transcription factor DAF-16/FOXO in the longevity mechanism of long-lived animals such as the Insulin-IGF-1 receptor *daf-2* mutants [18–20]. However, the tissue-specific role of key mechanisms in the proteostasis network, such as the 26S proteasome, remain incompletely understood. The 26S proteasome is a major degradation machinery of polyubiquitinated proteins [21], and aging *C. elegans* progressively accumulate proteasomal subunits, perhaps in a futile attempt to prevent further protein aggregation [22] that ultimately characterizes the age-related proteostatic collapse [23, 24]. Subunits in the 26S proteasome have been linked to lifespan determination [25, 26], but the tissue-specific contributions of proteasomal degradation in longevity are unclear and the tissue-specific dynamics of polyubiquitinated proteins is unknown.

Here, we systematically validated commonly used tissue-specific RNAi systems in *C. elegans* and found that the level of RNAi insensitivity of RNAi-defective strains and the level of specificity of tissue-specific RNAi strains unexpectedly vary widely. We uncovered that increasing temperature could improve RNAi insensitivity in some RNAi-defective mutants, potentially via further structural destabilization from point mutations. Leveraging validated tissue-specific RNAi strains, we identified the germline as a key tissue for proteasomal degradation, markedly overshadowing somatic tissues in proteasomal polyubiquitinated protein burden. We found that such burden is due in part by germline-acquired vitellogenins that are actively degraded during embryogenesis. We demonstrate the requirement for rigorous protein extraction protocols in the study of polyubiquitinated proteins during aging in *C. elegans* [27], including the use of aggregate-solubilizing buffers, and we highlight the need to study somatic and germline proteostasis separately in *C. elegans*. Finally, we uncover a novel role for the polyubiquitin gene *ubq-1* in the regulation of highly expressed genes including vitellogenins, suggesting a converging regulatory role for polyubiquitin in vitellogenin metabolism and lifespan.

## RESULTS

### RNAi-defectiveness of *rde-1* and *sid-1* mutants varies greatly and is modulated by temperature

Many RNAi-defective mutants (including *rde-3, rde-4, rde-10* and *rde-11*) retain RNAi sensitivity during short-term silencing, preventing their use to develop tissue-specific RNAi systems [13]. To unambiguously determine RNAi insensitivity, we tested the effect of whole-adulthood RNAi treatments in commonly used RNAi-defective mutants, including two *rde-1 (ne300* and *ne219)* [8] and two *sid-1 (qt9* and *pk3321)* mutants [28] that have established transgenic tissue-specific RNAi reconstitutions (**Supplementary Table S1**). To ensure that gene silencing provokes a remarkable effect on lifespan, we opted to silence *ubq-1,* a well-conserved gene (UBC in humans) encoding polyubiquitin [29]. Silencing *ubq-1* in wild-type N2 animals led to a marked mean lifespan decrease of about 50-60% at 20°C (**Figure 1A, Supplementary Table S2**). In different commonly used RNAi-defective mutants, silencing *ubq-1* at 20°C had surprisingly variable effects. Interestingly, the lifespan of *rde-1(ne300)* mutants was not affected by whole adulthood *ubq-1* silencing highlighting that they are indeed RNAi-insensitive (**Figure 1B**). Indeed, levels of *ubq-1* mRNA are not decreased in *rde-1(ne300)* mutants subjected to *ubq-1* RNAi (**Supplementary Figure S1A).** This is in line with previous observations of the lack of short-term RNAi effects when silencing fatty acid desaturases in *rde-1(ne300)* mutants [13], and confirms the functional abrogation of RDE-1 from this premature stop codon mutation. However, in *rde-1(ne219)* (point mutation)*, sid-1(qt9)* (premature stop codon) and *sid-1(pk3321)* (point mutation) mutants, *ubq-1* silencing resulted in significant deleterious effects on lifespan, indicating functionally hypomorphic variants that are not insensitive to RNAi at 20°C (**Figure 1C-E, Supplementary Table S2**). Since most studies in *C. elegans* are generally performed at 20°C, our data demonstrates that using RNAi-defective, but not fully RNAi-insensitive strains to study tissue-specific effects prevents appropriate interpretations.

**Figure 1.**
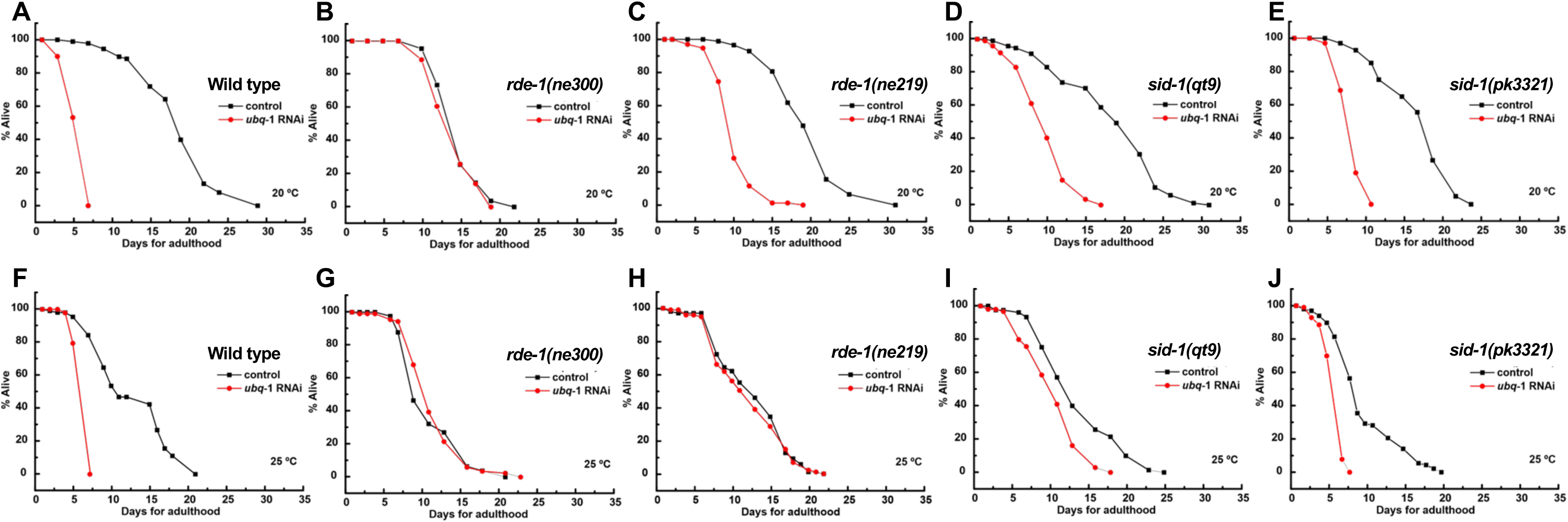
Levels of RNAi-defectiveness vary widely and in a temperature-dependent manner across *rde-1* and *sid-1* mutants. Lifespan of adult wild-type nematodes and the mutant strains *rde-1 (ne300) V*, *rde-1(ne219) V*, *sid-1 (qt9) V*, and *sid-1 (pk3321)* were analyzed at 20 °C **(A–E)** and 25 °C **(F–J)** and fed control bacteria or bacteria expressing dsRNA for *ubq-1*. Details on lifespan analyses are available in Supplementary Tables S2 and S3.

Since increasing temperature can affect the extent of the loss of function in mutants [30], we performed comparative lifespan analyses at 25°C in RNAi-defective mutants. Akin to 20°C, *ubq-1* silencing at 25°C resulted in a mean lifespan decrease of approximately 50% (**Figure 1F, Supplementary Tables S3**). This demonstrates that the capacity of the RNAi machinery to perform gene silencing was not affected by temperature, and the impact of silencing *ubq-1* on lifespan was correspondingly scaled between 20°C and 25°C [31, 32]. Expectedly, the *rde-1(ne300)* mutants remained insensitive to RNAi at 25°C (**Figure 1G, Supplementary Table S2**). Interestingly, silencing *ubq-1* in *rde-1(ne219)* mutants at 25°C did not affect their lifespan, indicating that the *rde-1(ne219)* point mutation becomes null at that temperature (**Figure 1H, Supplementary Table S2**). Since the hypomorphic loss of function at 20°C becomes null at 25°C in the *rde-1(ne219)* background, this strain could technically be used at 25°C for testing tissue-specific RNAi, resolving previous concerns for this strain in tissue-specific RNAi reconstitutions [13]. Increasing temperature to 25°C reduces the impact of *ubq-1* silencing on the lifespan of *sid-1(qt9)* and *sid-1(pk3321)*, suggesting that RNAi-sensitivity in these two mutants is also temperature-sensitive (**Figure 1I-J, Supplementary Table S2**). At both 20°C and at 25°C, *sid-1(qt9)* as well as in germline-less *sid-1(qt9);glp-1(e2144)* mutants remained sensitive to RNAi (**Supplementary Tables S2 and S3**). Moreover, *sid-1(pk3321)* mutants still retained significant RNAi sensitivity at 25°C (**Figure 1J**), corroborating a recent report highlighting this commonly used strain as incompatible for testing tissue-specific role of genes with RNAi [14]. Overall, our data indicates that only *rde-1(ne300)* is a suitable background at both 20°C and 25°C for interpreting tissue-specific RNAi strains, and we recommend experimenting at 25°C when using tissue-specific strains constructed from *rde-1(ne219)* mutants.

To investigate the structural impact of the *rde-1* and *sid-1* variants studied, we performed an *in silico* comparative analysis of the *rde-1* and *sid-1* proteins using UCSF ChimeraX [33]. Wild-type reference structures, obtained from the AlphaFold Database and the PDB (ID: 8XC1), were superimposed with mutant models generated via the AlphaFold server. and RMSD values were calculated exclusively over the structurally equivalent and conserved regions between the mutant proteins (*rde-1* and *sid-1*) and their respective wild-type counterparts (**Figure 2A-D**). The RMSD values relative to the wild-type protein, obtained for the selected atom pairs in *rde-1(ne300)* and *rde-1(ne219)*, were 0.954 Å and 0.845 Å, respectively. For *sid-1*, the RMSD values were 0.929 Å for *qt9* and 0.978 Å for *pk3321*. In the *rde-1(ne300)* allele, a premature stop codon in the PIWI domain leads to the loss of 197 amino acids (∼20% of the protein; **Figure 2A**) [8]. This region is essential for SID-1 activity, explaining the complete insensitivity of this strain to *ubq-1* RNAi-mediated gene silencing at both 20 and 25 °C in the lifespan assays. Conversely, the point mutation in the *rde-1(ne219)* allele results in the E414K substitution, replacing an acidic residue with a basic one in the PAZ domain (**Figure 2B**), a crucial region for siRNA recruitment and anchoring [34]. From a thermodynamic perspective, at 20 °C, the 3D structure of the PAZ domain retains sufficient conformational stability to sustain the silencing pathway. However, the increased kinetic energy at 25 °C potentially induces conditional instability, leading to local collapse or structural loosening. This interpretation converges with previous findings [34] demonstrating that the *ne219* allele compromises the global stability of the *rde-1* protein. Notably, in *sid-1(qt9)*, the transmembrane domain of SID-1 is absent (∼80% of the total protein; **Figure 2C**) [9, 28, 35], yet this strain remained partially sensitive to *ubq-1* silencing. In *sid-1(pk3321),* the charge alteration in the extracellular domain (D130N; **Figure 2D**) also only partially affect silencing sensitivity. In point mutations, increased temperature likely further destabilizes those key functional regions, ultimately reducing RNAi sensitivity. Taken together, our lifespan analyses clearly demonstrate that proper validation of RNAi-defective strains in the development of tissue-specific RNAi systems is essential. By increasing temperature to 25°C, we also uncovered that RNAi insensitivity can be fully attained in *rde-1(ne219)* mutants and can be partially improved in *sid-1(qt9)* and *sid-1(pk3321)* mutants, likely via functionally relevant structural destabilization.

**Figure 2.**
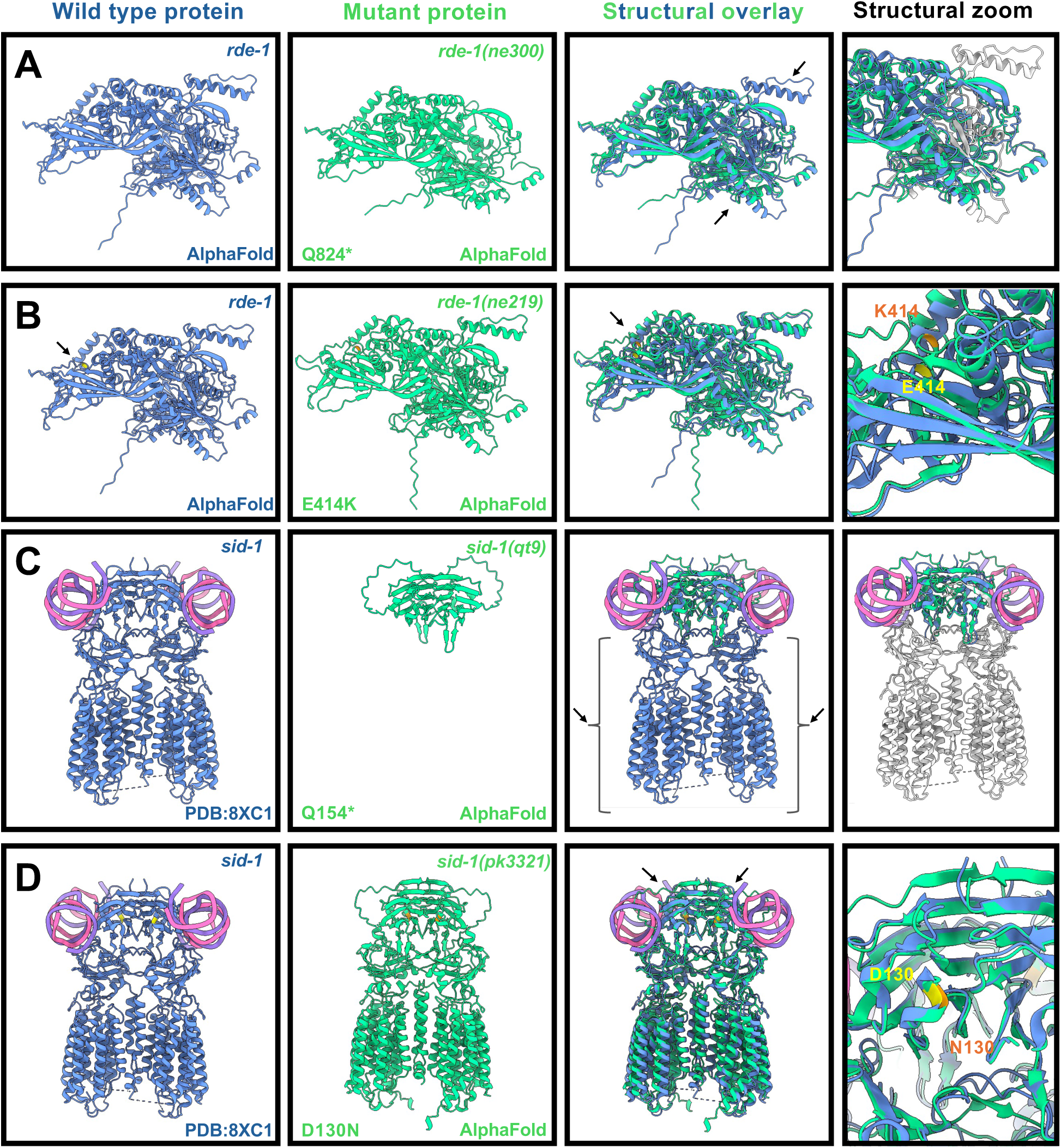
*In silico* structural analysis of *rde-1* and *sid-1* variants in *C. elegans*. **(A)** Impact of the ne300 allele on the *rde-1* protein (Q824*, green; AlphaFold server) relative to the wild-type protein (blue; AlphaFold server). The overlay highlights the loss of ∼20% of the structure in the Piwi domain; the missing region (197 amino acids) is represented in gray on the reference structure. **(B)** Structural superposition between wild-type *rde-1* (blue; AlphaFold Database) and the *ne219*-associated E414K variant (green; AlphaFold server). While the global architecture remains highly conserved, the focused view (right) illustrates the local environment change where the acidic glutamic acid (E414, yellow) is replaced by the basic lysine (K414, orange). **(C)** Impact of the qt9 allele on the *sid-1* protein (Q154*). The truncated form (green; AlphaFold Server) is overlaid on the wild-type structure (blue; PDB 8XC1), where the gray color delimits the loss of ∼ 80% of the protein (523 amino acids), including the transmembrane domain (TD). **(D)** Architecture of the *sid-1(pk3321)* protein. The model depicts the D130N substitution (green; AlphaFold server) aligned against the wild-type reference (blue; PDB: 8XC1). The magnified inset highlights the shift at position 130, contrasting the wild-type aspartic acid (D130, yellow) with the mutant asparagine (N130, orange). In the *sid-1* structure (8XC1), the dsRNA strands are highlighted in pink and purple. All molecular renderings and alignments were performed using UCSF ChimeraX.

### Tissue-specific RNAi strains are not always specific to their corresponding tissue

Despite numerous publications, authentic specificity of commonly used tissue-specific RNAi strains has never been fully characterized. Thus, we tested the specificity of previously engineered tissue-specific RNAi strains in both *rde-1* and *sid-1* background [13, 36–42] (**Supplementary Table S1**). To test the level of tissue-specificity, we leveraged important tissue-specific genes that create unequivocal morphological or behavioral changes in 100% of wild-type animals when silenced during development. For the intestine, we silenced *elt-2*, a transcription factor involved intestinal development and differentiation [43]. Loss of *elt-2* leads to underdeveloped and clear larvae. For the hypodermis, we silenced *bli-3*, an NADPH oxidase essential for molting, whereas its loss leads to a short animal with blisters [44]. For the muscle, we silenced *unc-54*, a myosin gene necessary for muscle function and mobility [45]. For the germline, we silenced *xpo-1,* a nuclear exportin important in germline development and lifespan [46, 47]. Low *xpo-1* during development affects the reproductive system and leads to curled adult animals and vulvar protrusion [46]. We silenced these genes in commonly used tissue-specific RNAi systems generally available at the Caenorhabditis Genetics Center (CGC) (**Supplementary Table S1**). Importantly, since the silencing was done on a short period of time (i.e. eggs to day 1 of adulthood), unlike our adult-only silencing lifespan analyses (**Figure 1A-J**), we did not observe any deleterious effect of tissue-specific RNAis in RNAi-defective strains, indicating that short periods of silencing may not suffice to phenotypically observe RNAi sensitivity in those strains (**Figure 3A-D**). We found that in *rde-1(ne300)* background [13], every tissue-specific RNAi strain behaved as advertised (**Figure 3A-D**). Notably, silencing *elt-2* in strains with intestinal RNAi sensitivity reproducibly generally resulted in 100% of animals displaying clear developmental deficits. In the *rde-1(ne219)* background, the annotated intestine-specific RNAi strain was also sensitive, albeit partially, to hypodermal gene silencing (**Figure 3A-D**). Although unclear, different extents of phenotypic representation could be due to varying levels of exposure of bacteria expressing dsRNA on agar plates. The annotated hypodermal-specific RNAi strain was also sensitive to intestinal and germline-related gene silencing (**Figure 3A-D**). The muscle-specific RNAi strain had RNAi sensitivity in muscle, but with relatively low silencing capacity (**Figure 3A-D**). The germline-intestine-specific and the germline-specific strains were limited to their annotated tissues (**Figure 3A-D**).

**Figure 3.**
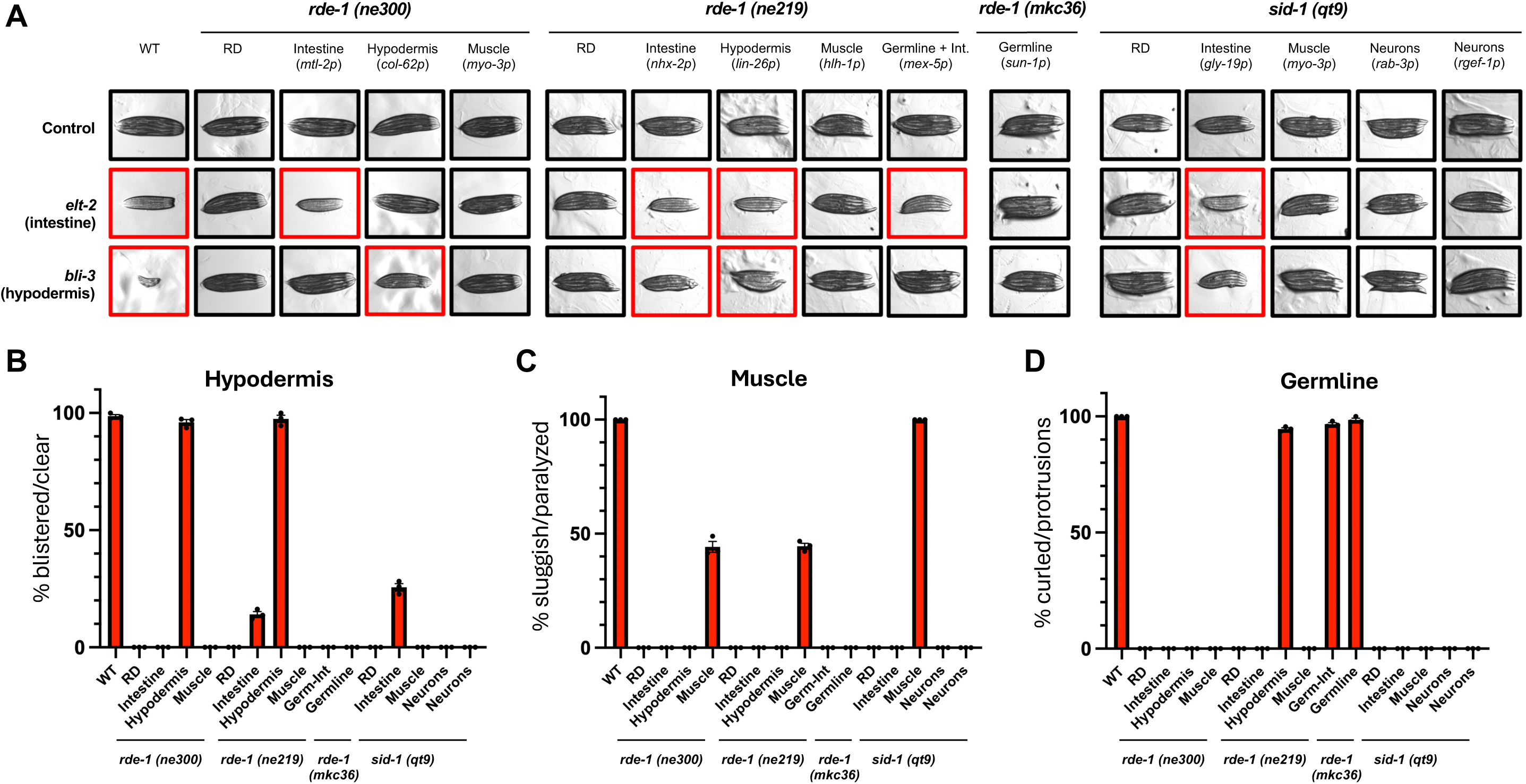
Tissue-specific RNAi systems display varying degrees of specificity. **(A)** WT, *rde-1* and *sid-1* mutants as well as different strains with annotated tissue-specific reconstitution for the intestine, hypodermis, muscle, germline or neurons were fed control bacteria or bacteria expressing dsRNA against *elt-2* (intestine), *bli-3* (hypodermis)*, unc-54* (muscle) or *xpo-1* (germline) from eggs to Day 1 of adulthood. In the case of partial effects with *bli-3* silencing, animals that showed blisters were presented. Quantification of displayed blisters (*bli-3* silencing) **(B)**, sluggishness (*unc-54* silencing) **(C)**, and vulvar protrusions (*xpo-1* silencing) **(D)**. Three independent experiments (n=33-57 per experiments).

In the *sid-1(qt9)* background, some tissue-specific reconstitutions also include a co-expressed fluorescent marker driven by the corresponding promoter to help visualize tissue-specificity (**Supplementary Table S1**). However, as observed with the *mtl-2* promoter in the *rde-1(ne219)* background, the intestine-specific RNAi strain designed with the *gly-19* promoter displayed partial RNAi sensitivity to *bli-3* silencing (**Figure 3A-D**). On the other hand, the muscle-specific strain was only sensitive in the muscle (**Figure 3A-D**). Neuronal-specific strains were not RNAi sensitive in peripheral tissues (**Figure 3A-D**), which differs from the strains originally built from *sid-1(pk3321)* [14]. Taken together, our analysis of tissue-specificity clearly lays out the limitations associated with reconstituting gene function under a particular promoter and highlights the importance of proper validation of commonly used tissue-specific RNAi-sensitivity reconstitutions prior to experimentation.

### Proteasomal degradation is most active in the germline

One of the leading questions in *C. elegans* aging studies is: how does the proteostasis network changes during aging? Longitudinal proteomic analyses in different nematodes have suggested that subunits of the 26S proteasome accumulate during aging [22], but their impact on overall polyubiquitinated protein metabolism is unclear. The 26S proteasome, which consists of a 19S cap and 20S proteolytic core, carries out the degradation of most polyubiquitinated proteins. Unexpectedly, in short-lived proteostatically-deficient *C.* elegans, including *hlh-30(tm1978)* [48], *hsf-1(sy441)* [49] and *daf-16(mu86)* [50], total levels of polyubiquitinated proteins did not significantly change at day 1 of adulthood (**Supplementary Figure S1B**). In wild-type *C. elegans*, we observed that the levels of polyubiquitinated proteins remains elevated during aging, using a complete solubilization protocol to avoid missing age-related protein aggregates (**Supplementary Figure S1C-F**). Thus, age-related accumulation of 26S proteasomes [22] does not seem to markedly affect overall levels of polyubiquitinated proteins. Seemingly, the steady-state total polyubiquitinated protein levels may not be a correlate of the overall status of the proteostasis network in different nematodes.

While many proteins are polyubiquitinated in *C. elegans*, the tissue-specific contribution of the total polyubiquitinated protein pool remains unknown. Using the established and broad tissue-specific RNAi system in the *rde-1(ne219)* background (**Supplementary Table S1**), we tested the role of the proteasome on polyubiquitinated protein accumulation by silencing 19S-related gene *rpn-6* and 20S-related gene *pas-7* [51] in tissue-specific strains validated in **Figure 2**. We also silenced *ubq-1* to better understand the role of polyubiquitin in different tissues. In wild-type animals, loss of proteasome led to a marked increase in total polyubiquitinated proteins, particularly in the higher molecular weight range, which was accompanied by increases in vitellogenin protein levels (**Figure 4A**, vitellogenins * and **) as observed by ponceau staining (antibodies for vitellogenins are unavailable). Six forms of vitellogenins exist, *vit-1* to *vit-6*, and they are the most abundant family of proteins in *C. elegans*, accounting for almost 20% of the total proteome [47]. While *vit-1* to *vit-5* are similar and overlap at 170kDa (*), *vit-6* is processed in two polypeptides of 115 and 88 kDa (**). Loss of *ubq-1* abolishes the presence of polyubiquitinated proteins, and surprisingly, led to a complete loss of vitellogenin proteins (**Figure 4A**). Conversely, RNAi-defective *rde-1(ne219)* mutants did not respond to any RNAi treatment (**Figure 4B**). In tissue-specific RNAi-sensitive reconstitution in the intestine and germline, we surprisingly saw limited increases in polyubiquitinated protein, suggesting a relatively limited role for the 26S proteasome in these tissues in the turnover of polyubiquitinated proteins (**Figure 4C**). In tissue-specific RNAi reconstitution in intestine, hypodermis and germline or intestine and germline, we observed similar vitellogenin and polyubiquitinated protein accumulation found in wild-type animals (**Figure 4D-E**), suggesting that the germline may be an important proteasomal degradation site. To directly confirm this, we silenced proteasome-related genes in a germline-specific RNAi reconstitution and found that limiting silencing to the germline is sufficient to accumulate polyubiquitinated proteins to a similar level than wild-type animals (**Figure 4F**). This demonstrates the unique and important role of proteasomal function in the germline and puts into context previous use of total polyubiquitinated protein levels as a proxy for organismal proteostasis. Here, we find that the extent of germline-specific and somatic proteasomal function markedly differs, a previously unrecognized difference that reveals important limitations in globally and indiscriminately studying polyubiquitinated protein metabolism and dynamics while inhibiting the proteasome systematically.

**Figure 4.**
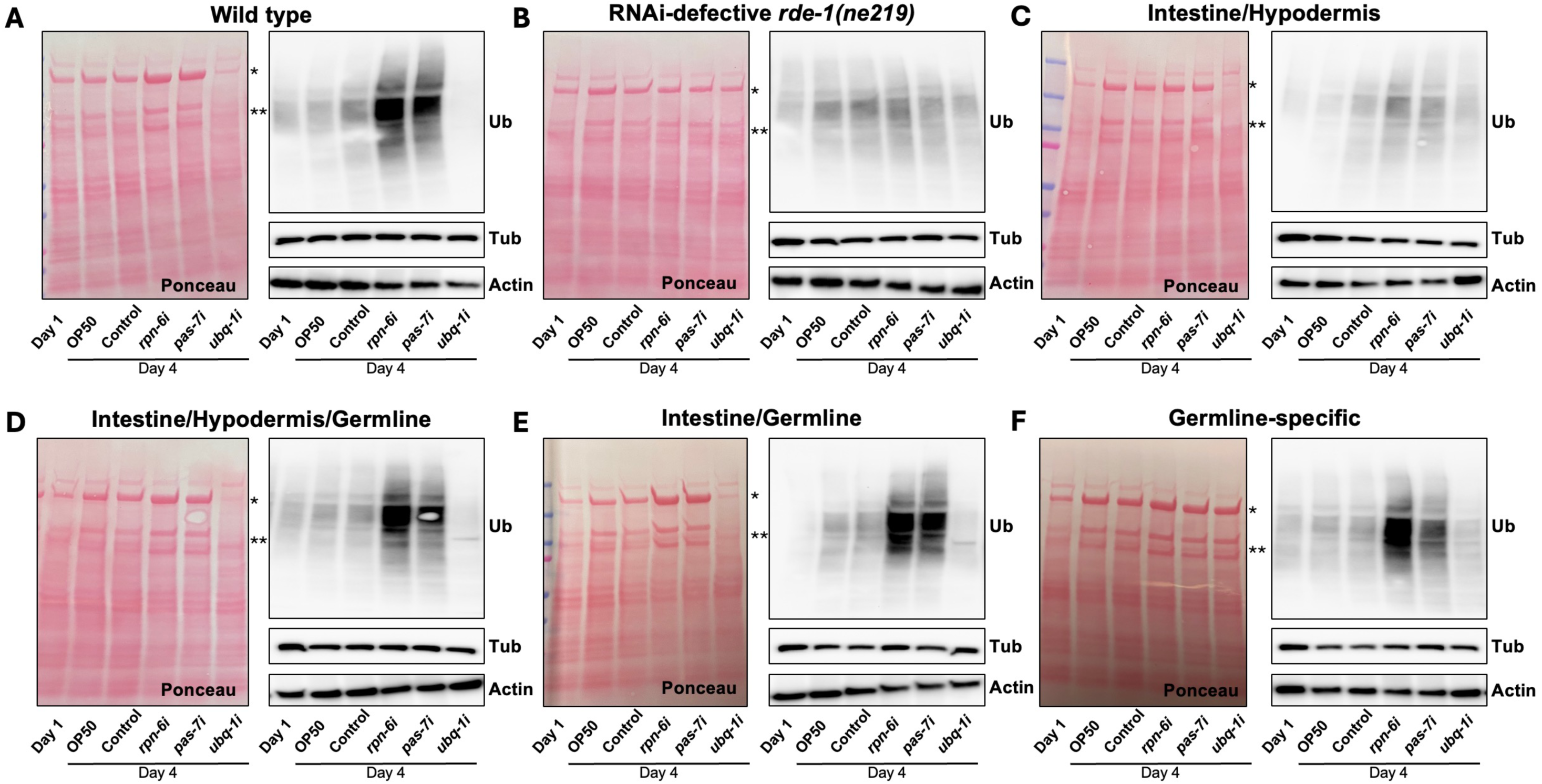
The germline is the most active site of proteasomal degradation in *C. elegans*. Adult-only gene silencing of *rpn-6, pas-7* and *ubq-1* from day 1 to day 4 of adulthood were performed at 25 °C in wild-type (**A**), *rde-1(ne219*) (**B**) as well as RDE-1 reconstituted strains in the intestine/germline (*nhx-2p*) (**C**), intestine/hypodermis/germline (*lin-26p*) (**D**), intestine/germline (*mex-5p*) (**E**), and germline (*sun-1p*) (**F**). Polyubiquitinated proteins (Ub) were visualized by immunoblotting and vitellogenin proteins (* and **) are highlighted in the adjacent Ponceau staining. Representative immunoblots of two independent experiments. Loading controls include immunoblotting for tubulin (Tub) and actin.

### Vitellogenin proteins in the germline are important contributors of total polyubiquitinated proteins

The inhibition of proteasomal function in the germline increased the levels of polyubiquitinated proteins and vitellogenins, while inhibition in the intestine and hypodermis had minimal impact on polyubiquitinated protein accumulation (**Figure 4D-F**). Thus, we reasoned that vitellogenin proteins, synthesized in the intestine and secreted thereafter, may be readily subjected to polyubiquitination only after entering the gonad, ultimately providing amino acids and lipids to oocytes via delipidation and proteasomal degradation. Here, we assessed the levels of total polyubiquitinated proteins at day 1 of adulthood in wild-type animals, *fer-15(b26);fem-1(hc17)* mutants that are sterile at 25°C and *glp-1(e2144)* mutants that are devoid of a germline at 25°C. Compared to wild-type animals, *fer-15;fem-1* mutants accumulated more polyubiquitinated proteins, which may suggest a tendency for these sterile animals to trap larger amounts of vitellogenin proteins in the unfertilized oocytes (**Figure 5A**). Conversely, since they lack a germline, *glp-1* mutants do not accumulate vitellogenin in the reproductive tissue due to its absence and of low expression of the vitellogenin endocytosis receptor *rme-2* [52, 53]. Germline-less animals carry significantly less polyubiquitinated proteins, despite having elevated vitellogenin accumulation, which is largely pseudocoelemic [53, 54] (**Figure 5A**). Since both sterile *fer-1;fem-15* and germline-less *glp-1* animals have dysfunctional reproductive systems, the levels of polyubiquitinated proteins remain largely the same later in adulthood (**Figure 5B**). Unexpectedly, silencing proteasomal subunits in *glp-1* animals did not result in overly large accumulation of polyubiquitinated proteins (**Figure 5C**), as seen in wild-type animals (**Figure 4A**). This observation combined with our data in intestine-hypodermis RNAi sensitive strain (**Figure 4C**) is rather perplexing considering the previously highlighted role for proteasomal function in the intestine of long-lived *glp-1* animals [25], and suggest relatively limited burden of the 26S proteasome in the soma compared to the germline. Using animals expressing VIT-2::GFP [52], we found that silencing *rpn-6* led to marked accumulation in the reproductive system while silencing *ubq-1* abrogated VIT-2::GFP levels (**Figure 5D**), corroborating our observations by ponceau (**Figure 4A**).

**Figure 5.**
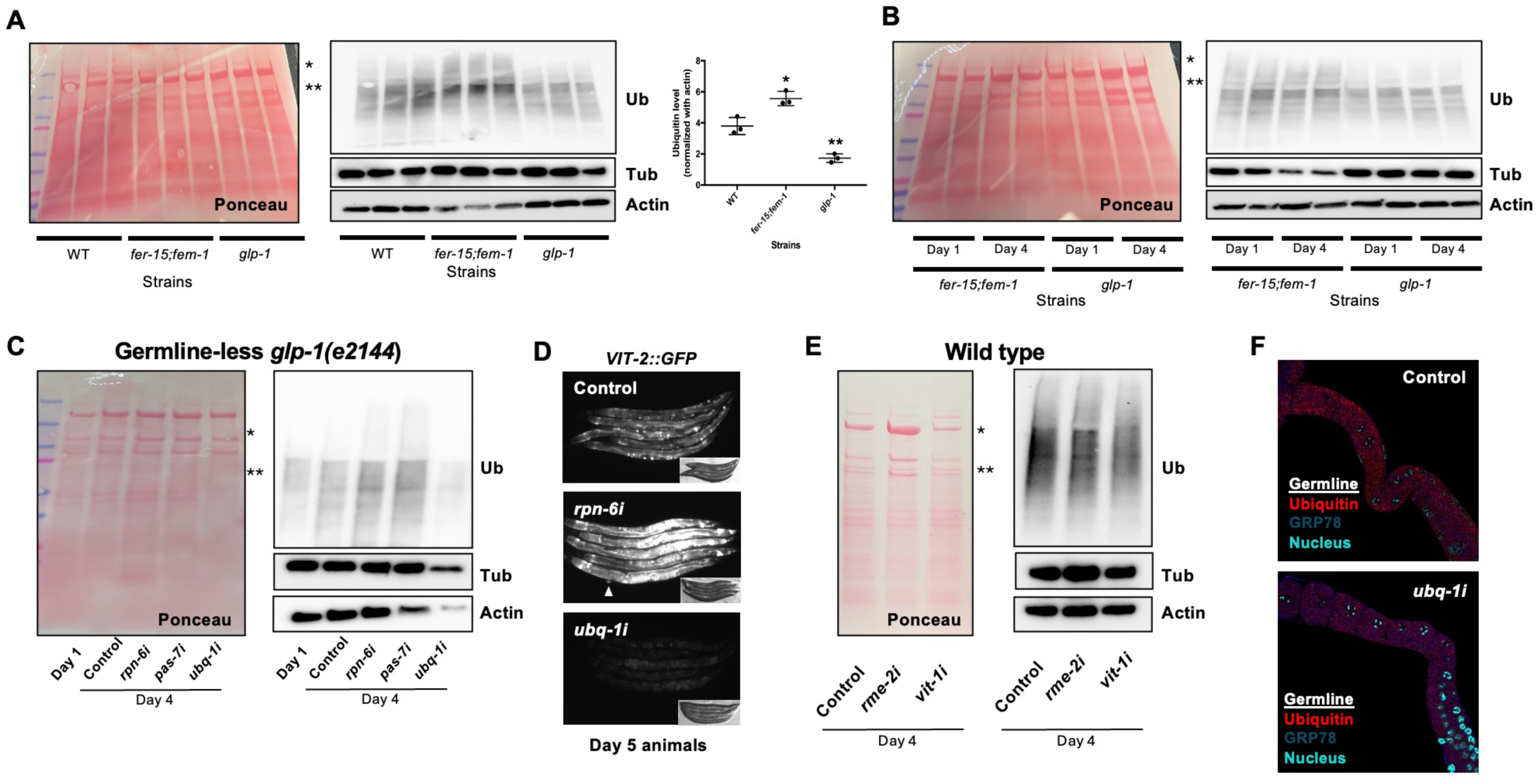
Vitellogenins located in the germline are major contributors of overall polyubiquitinated protein levels. **(A)** Polyubiquitinated protein levels were visualized by immunoblotting in Day 1 wild-type (WT), *fer-15(b26);fem-1(hc17)* and *glp-1(e2144)* animals raised at 25 °C on OP50 *E. coli*. Three independent replicates SD +/- *ANOVA *: P<0.05, **P<0.01*. (**B)** Polyubiquitinated protein levels were visualized by immunoblotting in day 1 and day 4 wild-type (WT) and *glp-1* animals raised at 25 °C on OP50 *E. coli*. (**C)** Adult-only silencing of *rpn-6, pas-7* and *ubq-1* in *glp-1* mutants raised at 25°C on OP50 *E. coli*. (**D)** Adult-only silencing of *rpn-6* and *ubq-1* from Day 1 to Day 5 in VIT-2::GFP expressing animals. (**E)** Adult-only silencing of *rme-2* and *vit-1* from Day 1 to Day 4 in wild-type animals. (**F)** Extracted germline from Day 3 wild-type animals fed control bacteria or bacteria expressing dsRNA against *ubq-1* from Day 1 of adulthood were visualized by immunofluorescence using antibodies against ubiquitin, ER-resident GRP78 and DAPI (nucleus).

To directly determine the role of germline-located vitellogenins in the total polyubiquitinated pool, we silenced the vitellogenin endocytosis receptor *rme-2* [52] as well as vitellogenins using *vit-1* RNAi as it efficiently silences all vitellogenins [53]. Lower total polyubiquitinated proteins were observed when transport of vitellogenins to the gonad is abrogated or when vitellogenins are silenced (**Figure 5E**). This is in line with previous proteomic analyses demonstrating the presence and accumulation of ubiquitinated vitellogenins in *C. elegans* [55]. Finally, in extracted gonadal tissue from wild-type animals, we observed ubiquitin-rich cytoplasmic clusters in developing oocytes, suggesting that the germline contains ample cargos, including vitellogenins, for proteasomal degradation (**Figure 5F**). This is supported by the presence of noticeably bright eggs layed by VIT-2::GFP expressing animals when proteasomal subunit *rpn-6* is inhibited (**Figure 5D**, arrow). Taken together, our data support a key role for the proteasome in vitellogenin metabolism in the germline, and an important contribution for gonadally located vitellogenins in overall polyubiquitinated protein levels in *C. elegans*.

### UBQ-1 is essential for the transcription of highly expressed genes

Our tissue-specific strain analysis of proteasome function inadvertently revealed a role for UBQ-1 in vitellogenin production (**Figure 4A-F**). Indeed, *ubq-1* silencing led to an almost complete abrogation of vitellogenin in animals sensitive for RNAi in the intestine (**Figure 4A, C-E**) while having no effect in the germline-specific RNAi strain (**Figure 4F**). Vitellogenins are the most abundant proteins and among the most expressed genes in *C. elegans* [47] and since they generally are secreted after being synthesized in the intestine, we reasoned that the significant reduction in vitellogenins may be due to a marked decrease in synthesis accompanied by secretion of existing vitellogenins and subsequent loss when eggs were laid. In addition to ponceau (**Figure 6A**), we observed the loss of vitellogenin with a VIT-2::GFP reporter when *ubq-1* was silenced (**Figure 5D**).

**Figure 6.**
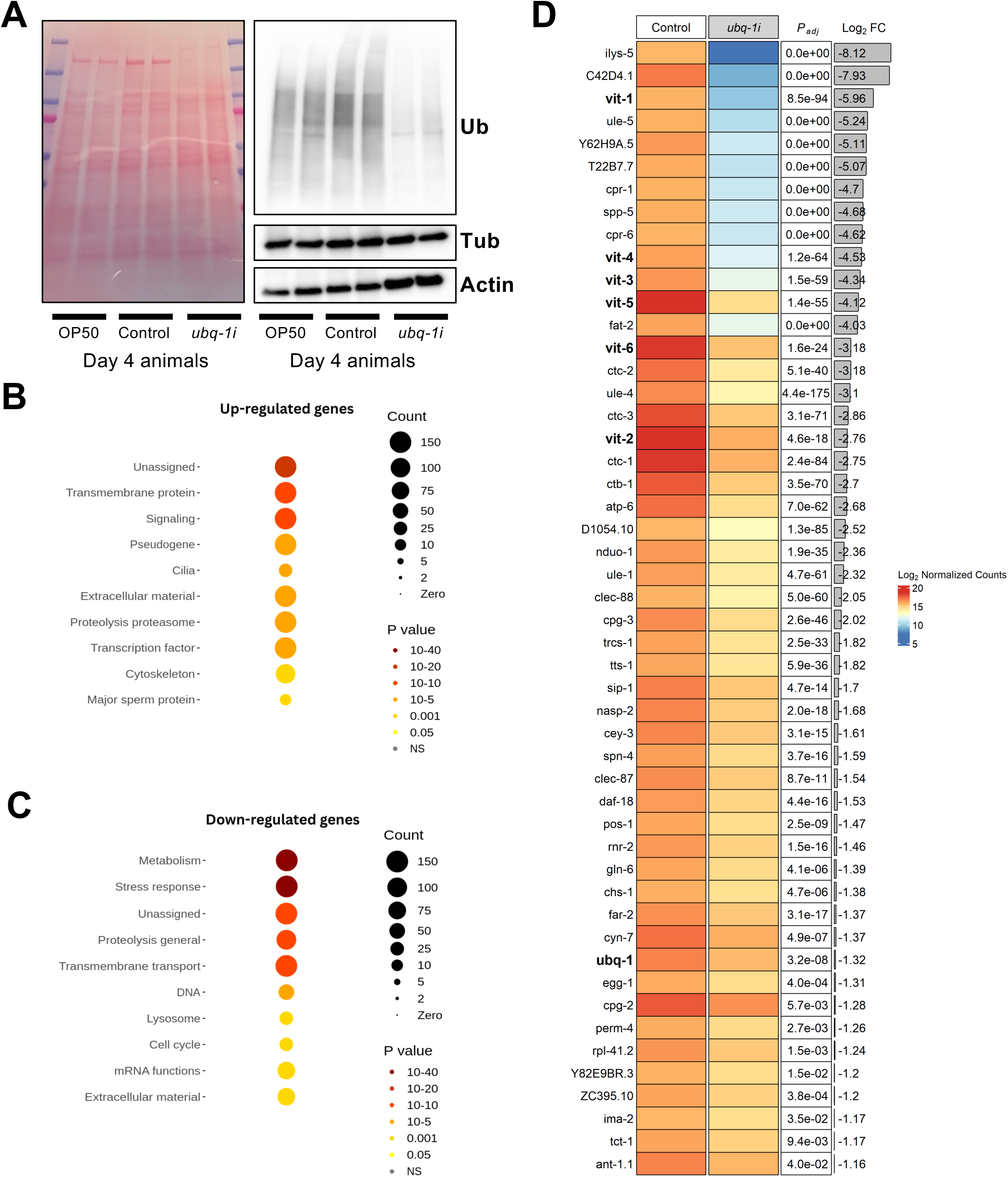
UBQ-1 is required for the active transcription of highly-expressed genes.(A) Polyubiquitinated protein levels in Day 4 wild-type animals fed OP50 *E.* coli, control bacteria or bacteria expressing dsRNA against *ubq-1.* Biological duplicates. (B) Wormcat analysis of upregulated genes and (C) downregulated genes in animals subjected to *ubq-1* silencing from Day 1 to Day 4 of adulthood (D) Top 50 highest expressed genes with significant downregulated expression from *ubq-1* silencing include marked downregulation of vitellogenin genes. Details on global expression changes are available in Supplementary Figure S2B and Supplementary Data S1.

To systematically understand the role of UBQ-1 in transcription, we silenced *ubq-1* during early adulthood and identified the global transcriptomic changes by RNA sequencing (**Supplementary Figure S2A-B, Supplementary Data S1**). Loss of *ubq-1* led to a drastic transcriptional rearrangement and pathway enrichment analysis suggested major downregulation of metabolic and stress response genes (**Figure 6B-C**). Interestingly, we found a massive reduction of many of the highest transcribed genes, with the largest downregulations featuring vitellogenin genes (**Figure 6D**). This indicated a key role for UBQ-1, not only in lifespan, but also in the active transcription of highly expressed genes. Taken together, our study demonstrates roles for the polyubiquitin gene UBQ-1 that extends beyond proteostasis to transcriptional specification and converges to modulate vitellogenin metabolism and lifespan.

## Discussion

Significant advances have been made over the last decade to develop tools for spatial and tissue-specific study of the role of genes in *C. elegans* biology. Unfortunately, a complete validation of available tissue-specific RNAi systems over the lifespan of *C. elegans* had yet to be performed. Even after several years and hundreds of publications, proper demonstration of RNAi insensitivity in RNAi-defective mutants remained an important validation of tissue-specific RNAi systems allowing proper studies of tissue-specific function of genes in *C. elegans*. Here, we thoroughly determined the validity of different RNAi-defective strains for the development and use of tissue-specific RNAi strains.

Leveraging the efficient lifespan-shortening effects of silencing polyubiquitin gene *ubq-1, w*e found that *rde-1(ne300)* is a robust strain beyond short-term silencing [13] and across the whole lifespan of *C. elegans*. Corresponding tissue-specific strains were also restricted for RNAi sensitivity to their specific tissue. We uncovered that the *rde-1(ne219)* are only fully insensitive to RNAi at 25°C, supporting a temperature-sensitive null mutation via local structural destabilization at the point mutation. Our whole adulthood silencing approach strengthen our understanding of RNAi-defective strains from short-term RNAi studies [34, 56] and provide unequivocal answers on the RNAi insensitivity necessary for the construction of tissue-specific RNAi system. Notably, tissue-specific RNAi strains in the *rde-1(ne219)* and *sid-1(qt9)* originally identified for the intestine or the hypodermis, but sensitive in either tissue or more, will need to be re-annotated accordingly. Finally, we conclude that the *sid-1(qt9)* and *sid-1(pk3321)* mutants are ill-suited for the development of tissue-specific RNAi strains, because they retain substantial long-term RNAi capacity at both 20°C and 25°C. While our tissue-specific validation of RNAi systems using short-term silencing suggest that tissue reconstitutions in *sid-1(qt9)* mutants are generally robust, studies using long-term silencing are not advisable in that background. Additionally, tissue-specific reconstitution of SID components was recently shown to negatively affect lifespan [57]. Notably, complete loss of *sid-1* may further improve RNAi insensitivity [58], but the effects observed in *sid-1 (qt9)* mutants, which express a severely truncated SID-1, suggest potentially non-negligible SID-1-independent dsRNA import in tissues. Therefore, the numerous studies using *sid-1(qt9)* and *sid-1(pk3321)* and the corresponding tissue-specific reconstitutions should be re-evaluated with our observations in mind. Altogether, our study systematically clarified the issues related to the development and use of tissue-specific strains in *C. elegans* and uncovered a solution (i.e. experiments carried out at 25°C) to address incomplete RNAi insensitivity in *rde-1(ne219)* mutants at 20°C.

The role of the 26S proteasome in the lifespan of *C. elegans* and the dynamics of polyubiquitinated proteins have been previously highlighted [25, 26, 55], but tissue-specific contributions were unclear and proper polyubiquitinated protein analyses that included aggregated proteins were lacking. Here, we uncovered an important role for the 26S proteasome in the germline, especially in the metabolism of polyubiquitinated vitellogenin proteins. Germline-specific silencing of 26S proteasome subunits led to similar accumulation of polyubiquitinated proteins to whole-organism silencing, demonstrating that whole-organism analyses of polyubiquitinated proteins is relatively misleading, especially when trying to study somatic maintenance and longevity. This is particularly evident when levels of polyubiquitinated proteins are analysed longitudinally (**Supplementary Figure S1C-F**) and comparatively in different short-lived, sterile and germline-less animals (**Figures 5A and Supplementary S1B**). Indeed, total levels of polyubiquitinated proteins in *C. elegans* are an incomplete proxy for the organismal-wide status of the proteostasis network.

In our study, we find that the key converging feature for polyubiquitinated protein accumulation in *C. elegans* is vitellogenin metabolism in the germline. As observed in germline-less animals that accumulate vitellogenin proteins in the pseudocoelom [53, 54], intestinally synthesized vitellogenins are likely only ubiquitinated when it reaches the gonads. Vitellogenin structure, homologous to the N-terminus of human apolipoprotein B (apoB), is a large lipid transfer protein with amphipathic beta-sheets that accommodate lipids to a high-density lipoprotein level [59]. The relatively compact structure of vitellogenins accompanied by co-translational lipidation in the endoplasmic reticulum generally protects nascent lipoproteins from co-translational ubiquitination observed with the significantly larger and more hydrophobic apoB protein [60]. However, endocytosis in the germline and transport to developing embryos likely destabilizes vitellogenins, fostering their ubiquitination [55] and consequent proteasomal degradation. In *Caurosius morosus* (stick insect) vitellin, a processed formed of vitellogenin, is ubiquitinated and degraded in the developing embryos [61]. We previously found that lipid-droplet enriched fraction contains high levels of polyubiquitinated proteins when *rpn-6* is silenced, and a portion of that accumulation may be related to germline-associated vitellogenins [62]. Thus, we conclude that vitellogenins are ubiquitinated in the germline, potentiated by destabilization as it enters the germline and as its lipids gets progressively mobilized by the developing embryos. The appearance of undegraded vitellogenin proteins in laid eggs when proteasomal function is inhibited supports this mechanism (**Figure 5D**).

Vitellogenin protein degradation dynamics in the germline could therefore largely account for variations in total polyubiquitinated protein levels in different *C. elegans* strains. In wild-type animals (and in short-lived animals), brood size is similar and brood production ends within the first 5 days of adulthood. In *fer-15;fem-1* sterile animals, the inability to generate viable progeny leads to increased accumulation of polyubiquitinated proteins in the germline. Proteostasis challenges associated with sterility have been previously reported [63]. The role of the 26S proteasome in the germline was also recently highlighted in the context of sex determination and transgenerational information transfer [64]. The rate of progeny production likely also plays a role in determining steady-state total polyubiquitinated protein levels. Reported vulvar venting of vitellogenins from adults to progenies could conceivably contribute to fluctuations in total polyubiquitinated proteins [65]. Overall, our data points to important tissue-specific variations in proteasomal burden, especially between the germline and the soma, and centers vitellogenins as important contributors to total observed polyubiquitinated proteins in *C. elegans*.

The polyubiquitin gene UBQ-1/UBC is important for normal lifespan in *C. elegans* [66], and its role in proteostasis is well established [67]. When expressed, this polyubiquitin protein gets cleaved by UBC hydrolases to form ubiquitin available for the ubiquitin-proteasome system [68]. Ubiquitin can also serve as a transcriptional activator or repressor when conjugated with histones, affecting chromatin compaction and promoter accessibility [69, 70]. Here, we found that loss of *ubq-1* generates important and broad transcriptional changes that likely contribute to the accelerated aging observed. Interestingly, despite being highly expressed throughout adulthood, vitellogenins are among the most downregulated genes from the loss of *ubq-1*, highlighting the major role of ubiquitin on highly active transcription. Because of the massive downregulation of those genes, ubiquitin may be essential for the transcriptional machinery to access those corresponding loci. Alternatively, ubiquitin-dependent chromatin dynamics on typically repressed gene loci may be lost potentially leading to re-positioning of transcription factors and machineries onto typically lowly-expressed genes, thereby minimizing the expression of otherwise highly expressed genes. As aging is characterized by progressive transcriptional drift due in part by age-related epigenetic changes [71, 72], our study positions the maintenance of UBQ-1 expression and function as a key proteostatic and transcriptional modulator in the health and lifespan of *C. elegans*.

Importantly, our study demonstrates the importance of proper initial validation of tools to study gene function, in particular tissue-specific RNAi systems. We conclude that the use of *rde-1* strains to construct tissue-specific strains is a superior approach to the use of *sid-1* strains. Here, we also provide an optimized protocol for the analysis of polyubiquitinated proteins, with a focus on complete homogenization to help reach whole proteome analyses and improve interpretations. Our original goal of authenticating tissue-specific RNAi strains led us to uncover major differences in proteasomal function between the soma and the germline, placing vitellogenins as important polyubiquitinated targets in developing embryos. Finally, we inadvertently uncovered that the polyubiquitin gene UBQ-1 is an essential protein in the proper maintenance of transcription, especially of highly expressed genes such as vitellogenins, placing it as a converging modulator of vitellogenin metabolism and lifespan.

## METHODS

### *C. elegans* maintenance, growth and visualization

Strains of *C. elegans* were grown and maintained at 20°C on growth media agar plates seeded with *E. coli* strain OP50 and handled as previously described [73]. Ethical approval was not required for nematode experiments by our institution. Strains used in this study are listed in **Supplementary Table S1**. For imaging, nematodes were immobilized with 0.1% sodium azide and visualized with a V20 fluorescence stereomicroscope (Zeiss). Exposure and magnification were maintained between conditions.

### Lifespan analyses and silencing experiments

Nematodes were synchronized by hypochlorite treatment and grown at 20 °C or 25 °C on *E. coli* OP50 until the first day of adulthood. Animals were then maintained at the same temperatures on plates seeded with either control bacteria (*E. coli* HT115 carrying the L4440 empty vector) or bacteria producing double-stranded RNA (dsRNA) targeting the *ubq-1* gene (Ahringer library) [2]. Animal survival was assessed every 2–3 days and tissue-specific silencing was performed using RNAi against *elt-2* (Ahringer library), *bli-3* (Vidal library), *unc-54* (Ahringer library), *xpo-1* (Ahringer library). Graphs were created using OriginPro 9 (OriginLab) and statistical analyses (Mantel-Cox log-rank) were performed using the OASIS 2 (https://sbi.postech.ac.kr/oasis2/surv/).

### Protein solubilization for immunoblotting

To completely solubilize proteins, we used a RIPA solution (50mM Tris-HCl, 150mM NaCl, 1mM EDTA, Roche Complete Protease Inhibitors) containing 5% SDS followed by heating for 5 minutes at 95□C and sonication for 20 seconds (QSonica Q55, 30% input). Insoluble proteins were pelleted by centrifugation for 2 minutes at 12000rpm. This first solubilization step (250□l of 5% SDS-RIPA) solubilized over 90% of total proteins, and a second step of solubilization (250□l of 5% SDS-RIPA) successfully solubilized the small pellet. Ubiquitinated proteins from combined (i.e. total) were visualized by immunoblotting. Mild homogenization (MH) was also performed using the buffer (250□l) and method described in Koyuncu et al. [55] and insoluble proteins were pelleted by centrifugation for 2 minutes at 12000rpm. Subsequently, these larger pellets (compared to the 5% SDS-RIPA method) were completely solubilized with 5% SDS-RIPA, heat and sonication as described above. Polyubiquitinated proteins were visualized by immunoblotting for ubiquitin (Millipore Sigma 05-994, clone P4D1-A11), and actin (Millipore Sigma MAB1501R) and tubulin (Abcam ab6160) was used as loading controls.

### RNA preparation and analysis

Approximately 3,000 nematodes from synchronized populations were cultured at 20 °C or 25 °C per independent replicate, with three or four biological replicates used for the qPCR and RNA-seq experiments. Animals were fed *E. coli* OP50 until reaching adulthood and then transferred to plates seeded with either control bacteria or bacteria expressing dsRNA, as previously described [47]. Adult nematodes were separated from the progeny by collection in M9 buffer followed by gravity sedimentation for 2 min, and were transferred daily to fresh RNAi plates until the final day of collection. After 48 or 72 hours of RNAi exposure, worms were collected with M9, pelleted, frozen at −80 °C, and used for RNA extraction. Total RNA was isolated using TRIzol reagent and subsequently purified with the Direct-zol RNA MiniPrep kit (Zymo Research). RNA concentration was determined by spectrophotometry using a TapeStation (Agilent). For qPCR analyses, 1 µg of RNA per sample was reverse transcribed into cDNA, and transcript levels were quantified as previously described [12] using a QuantStudio5 apparatus (Applied Biosystems). Primer sequences used for qPCR analyses in this study are listed in **Supplementary Table S4.** Statistical analyses were performed using GraphPad Prism 10 (GraphPad Software).

### RNA sequencing and differential gene expression analyses

Library construction and next-generation sequencing (NGS) were performed by polyadenylate selection and Illumina HiSeq methods (Genewiz-Azenta). Genes with a minimum of 10 read counts in 4 samples (the number of biological replicates per condition) were included in the analysis. The R package DESeq2 [74] was used to test genes for differential expression and those with a false discovery rate of < 0.05 and fold change > 2-fold were considered significant. GO enrichment analysis was conducted using WormCat 2.0 [75] and heatmaps of gene expression were created using the R package ComplexHeatmap [76].

### Germline extraction and immunofluorescence

For immunofluorescence staining, adult worms were dissected in 1x PBS on a coverslip. Coverslips were inverted onto poly-L-lysine–coated slides, snap-frozen at −80 °C, and subjected to freeze-crack. Slides were immersed in cold methanol for 1 min and fixed for 30 min in a humid chamber in 1x PBS, 0.08 M HEPES (pH 6.9), 1.6 mM MgSO₄, 0.8 mM EGTA, and 3.7% paraformaldehyde. Slides were washed three times in PBST (1x PBS, 0.1% Tween 20), blocked for 1 hr in PBST containing 3% BSA, and incubated overnight at 4 °C with primary antibodies diluted in PBST with 1% BSA. Primary antibodies used were anti-ubiquitin (Invitrogen, MA1-10035; 1:100), and anti-GRP78/HSPA5 (Novus Biologicals, NBP1-06274; 1:200). Secondary Alexa Fluor 555-conjugated goat anti-mouse (A32727, Life Technologies) and Alexa Fluor 647-conjugated goat anti-rabbit secondary antibodies (111-605-003, Jackson ImmunoResearch) at 1:200 were incubated for 1 hr at room temperature. Slides were washed in PBST, counterstained with DAPI, mounted in Vectashield, sealed, and stored at −20 °C until imaging on a FV3000 confocal microscope (Olympus).

### Computational analysis of RDE-1 and SID-1 structures

The three-dimensional structures of the *Caenorhabditis elegans* proteins *rde-1* and *sid-1* were obtained from public databases. The predicted structure of *rde-1* was retrieved from the Alphafold protein structure database (https://alphafold.ebi.ac.uk/), using the model AF-G5EEH0-F1-v6, corresponding to UniProt accession G5EEH0. The structure of *sid-1* was obtained from the Protein Data Bank (PDB), using 8XC1, corresponding to the dsRNA-bound form. Based on the amino acids sequence, point mutations were introduced using AlphaFold Server (https://alphafoldserver.com/). The mutants *rde-1 (ne300)* (Q824*)*, rde-1(ne219)* (E414K), *sid-1 (qt9)* (Q154*) *sid-1(pk3321)* (D130N) were generated, and native and mutant structures were subsequently visualized and analyzed using UCSF ChimeraX (https://www.cgl.ucsf.edu/chimerax/).

## ACKNOWLEDGEMENTS

We would like to thank Dr. Brad Morrison (Boise State University), Dr. Eric Allain and Sophie Landry (Université de Moncton), as well as Karine Blais for their feedback. We are grateful to the Malene Hansen laboratory (Buck Institute for Research on Aging) for sharing the N2 wild-type, MAH85, MAH677, AGD 635, AGD 637 and AGD639 used in this study. We would like to thank Dr. Patrick Narbonne (UQTR) for helping perform tissue extraction and immunofluorescence analyses. NSMDS performed lifespan analyses, structural analyses and mRNA extractions. CB performed lifespan analyses and tissue-specific RNAi studies. AO performed germline extraction and analyses. LH analyzed RNA sequencing data. SK performed lifespan analyses. SD and STD conducted longitudinal polyubiquitinated protein analyses. LRL designed the study, performed protein and gene expression analyses, and tissue-specific RNAi analyses, and wrote the manuscript. All co-authors edited the manuscript. No AI-assisted technologies were used in the writing of this manuscript. This work was funded by a Natural Sciences and Engineering Research Council of Canada (NSERC) Discovery Grant, a Project Grant from Canadian Institutes of Health Research (CIHR 519531) and a Research Chair in Precision Medicine (J.-Louis Lévesque Foundation – NB Research) to LRL.

## REFERENCES

1. Fire, A., et al., Potent and specific genetic interference by double-stranded RNA in Caenorhabditis elegans. Nature, 1998. 391(6669): p. 806–11.

2. Kamath, R.S., et al., Systematic functional analysis of the Caenorhabditis elegans genome using RNAi. Nature, 2003. 421(6920): p. 231–7.

3. Rual, J.F., et al., Toward improving Caenorhabditis elegans phenome mapping with an ORFeome-based RNAi library. Genome Res, 2004. 14(10B): p. 2162–8.

4. Hansen, M., et al., New genes tied to endocrine, metabolic, and dietary regulation of lifespan from a Caenorhabditis elegans genomic RNAi screen. PLoS Genet, 2005. 1(1): p. 119–28.

5. Hamilton, B., et al., A systematic RNAi screen for longevity genes in C. elegans. Genes Dev, 2005. 19(13): p. 1544–55.

6. Kenyon, C.J., The genetics of ageing. Nature, 2010. 464(7288): p. 504–12.

7. Lapierre, L.R. and M. Hansen, Lessons from C. elegans: signaling pathways for longevity. Trends Endocrinol Metab, 2012. 23(12): p. 637–44.

8. Tabara, H., et al., The rde-1 gene, RNA interference, and transposon silencing in C. elegans. Cell, 1999. 99(2): p. 123–32.

9. Winston, W.M., C. Molodowitch, and C.P. Hunter, Systemic RNAi in C. elegans requires the putative transmembrane protein SID-1. Science, 2002. 295(5564): p. 2456–9.

10. Zhang, J., et al., Structural insights into double-stranded RNA recognition and transport by SID-1. Nat Struct Mol Biol, 2024. 31(7): p. 1095–1104.

11. Stefanakis, N., I. Carrera, and O. Hobert, Regulatory Logic of Pan-Neuronal Gene Expression in C. elegans. Neuron, 2015. 87(4): p. 733–50.

12. Wong, S.Q., et al., Neuronal HLH-30/TFEB modulates peripheral mitochondrial fragmentation to improve thermoresistance in Caenorhabditis elegans. Aging Cell, 2023. 22(3): p. e13741.

13. Watts, J.S., et al., New Strains for Tissue-Specific RNAi Studies in Caenorhabditis elegans. G3 (Bethesda), 2020. 10(11): p. 4167–4176.

14. Gahlot, S. and J. Singh, Caenorhabditis elegans neuronal RNAi does not render other tissues refractory to RNAi. Proc Natl Acad Sci U S A, 2024. 121(22): p. e2401096121.

15. Kumsta, C. and M. Hansen, C. elegans rrf-1 mutations maintain RNAi efficiency in the soma in addition to the germline. PLoS One, 2012. 7(5): p. e35428.

16. Calixto, A., et al., Enhanced neuronal RNAi in C. elegans using SID-1. Nat Methods, 2010. 7(7): p. 554–9.

17. Zhang, L., et al., The auxin-inducible degradation (AID) system enables versatile conditional protein depletion in C. elegans. Development, 2015. 142(24): p. 4374–84.

18. Wolkow, C.A., et al., Regulation of C. elegans life-span by insulinlike signaling in the nervous system. Science, 2000. 290(5489): p. 147–50.

19. Libina, N., J.R. Berman, and C. Kenyon, Tissue-specific activities of C. elegans DAF-16 in the regulation of lifespan. Cell, 2003. 115(4): p. 489–502.

20. Zhang, Y.P., et al., Intestine-specific removal of DAF-2 nearly doubles lifespan in Caenorhabditis elegans with little fitness cost. Nat Commun, 2022. 13(1): p. 6339.

21. Collins, G.A. and A.L. Goldberg, The Logic of the 26S Proteasome. Cell, 2017. 169(5): p. 792–806.

22. Walther, D.M., et al., Widespread Proteome Remodeling and Aggregation in Aging C. elegans. Cell, 2015. 161(4): p. 919–32.

23. Ben-Zvi, A., E.A. Miller, and R.I. Morimoto, Collapse of proteostasis represents an early molecular event in Caenorhabditis elegans aging. Proc Natl Acad Sci U S A, 2009. 106(35): p. 14914–9.

24. Vecchi, G., et al., Proteome-wide observation of the phenomenon of life on the edge of solubility. Proc Natl Acad Sci U S A, 2020. 117(2): p. 1015–1020.

25. Vilchez, D., et al., RPN-6 determines C. elegans longevity under proteotoxic stress conditions. Nature, 2012. 489(7415): p. 263–8.

26. Keith, S.A., et al., Graded Proteasome Dysfunction in Caenorhabditis elegans Activates an Adaptive Response Involving the Conserved SKN-1 and ELT-2 Transcription Factors and the Autophagy-Lysosome Pathway. PLoS Genet, 2016. 12(2): p. e1005823.

27. Mills, J., et al., Combined Nucleotide and Protein Extractions in Caenorhabditis elegans. J Vis Exp, 2019(145).

28. Whangbo, J.S., et al., SID-1 Domains Important for dsRNA Import in Caenorhabditis elegans. G3 (Bethesda), 2017. 7(12): p. 3887–3899.

29. Stringham, E.G., D. Jones, and E.P. Candido, Expression of the polyubiquitin-encoding gene (ubq-1) in transgenic Caenorhabditis elegans. Gene, 1992. 113(2): p. 165–73.

30. Hirsh, D. and R. Vanderslice, Temperature-sensitive developmental mutants of Caenorhabditis elegans. Dev Biol, 1976. 49(1): p. 220–35.

31. Miller, H., et al., Genetic interaction with temperature is an important determinant of nematode longevity. Aging Cell, 2017. 16(6): p. 1425–1429.

32. Stroustrup, N., et al., The temporal scaling of Caenorhabditis elegans ageing. Nature, 2016. 530(7588): p. 103–7.

33. Goddard, T.D., et al., UCSF ChimeraX: Meeting modern challenges in visualization and analysis. Protein Sci, 2018. 27(1): p. 14–25.

34. Waddell, B.M. and C.W. Wu, Mutant alleles of the Caenorhabditis elegans rde-1 gene identified through chemical mutagenesis of an snRNA misprocessing reporter. G3 (Bethesda), 2025. 15(7).

35. Wang, R., et al., Structural basis for double-stranded RNA recognition by SID1. Nucleic Acids Res, 2024. 52(11): p. 6718–6727.

36. Yigit, E., et al., Analysis of the C. elegans Argonaute family reveals that distinct Argonautes act sequentially during RNAi. Cell, 2006. 127(4): p. 747–57.

37. Espelt, M.V., et al., Oscillatory Ca2+ signaling in the isolated Caenorhabditis elegans intestine: role of the inositol-1,4,5-trisphosphate receptor and phospholipases C beta and gamma. J Gen Physiol, 2005. 126(4): p. 379–92.

38. Qadota, H., et al., Establishment of a tissue-specific RNAi system in C. elegans. Gene, 2007. 400(1-2): p. 166–73.

39. Marre, J., E.C. Traver, and A.M. Jose, Extracellular RNA is transported from one generation to the next in Caenorhabditis elegans. Proc Natl Acad Sci U S A, 2016. 113(44): p. 12496–12501.

40. Zou, L., et al., Construction of a germline-specific RNAi tool in C. elegans. Sci Rep, 2019. 9(1): p. 2354.

41. Liu, L., C. Ruediger, and M. Shapira, Integration of Stress Signaling in Caenorhabditis elegans Through Cell-Nonautonomous Contributions of the JNK Homolog KGB-1. Genetics, 2018. 210(4): p. 1317–1328.

42. Gelino, S., et al., Intestinal Autophagy Improves Healthspan and Longevity in C. elegans during Dietary Restriction. PLoS Genet, 2016. 12(7): p. e1006135.

43. McGhee, J.D., et al., ELT-2 is the predominant transcription factor controlling differentiation and function of the C. elegans intestine, from embryo to adult. Dev Biol, 2009. 327(2): p. 551–65.

44. Edens, W.A., et al., Tyrosine cross-linking of extracellular matrix is catalyzed by Duox, a multidomain oxidase/peroxidase with homology to the phagocyte oxidase subunit gp91phox. J Cell Biol, 2001. 154(4): p. 879–91.

45. Moerman, D.G., et al., Mutations in the unc-54 myosin heavy chain gene of Caenorhabditis elegans that alter contractility but not muscle structure. Cell, 1982. 29(3): p. 773–81.

46. Silvestrini, M.J., et al., Nuclear Export Inhibition Enhances HLH-30/TFEB Activity, Autophagy, and Lifespan. Cell Rep, 2018. 23(7): p. 1915–1921.

47. Kumar, A.V., et al., Exportin 1 modulates life span by regulating nucleolar dynamics via the autophagy protein LGG-1/GABARAP. Sci Adv, 2022. 8(13): p. eabj1604.

48. Lapierre, L.R., et al., The TFEB orthologue HLH-30 regulates autophagy and modulates longevity in Caenorhabditis elegans. Nat Commun, 2013. 4: p. 2267.

49. Hsu, A.L., C.T. Murphy, and C. Kenyon, Regulation of aging and age-related disease by DAF-16 and heat-shock factor. Science, 2003. 300(5622): p. 1142–5.

50. Kenyon, C., et al., A C. elegans mutant that lives twice as long as wild type. Nature, 1993. 366(6454): p. 461–4.

51. Davy, A., et al., A protein-protein interaction map of the Caenorhabditis elegans 26S proteasome. EMBO Rep, 2001. 2(9): p. 821–8.

52. Grant, B. and D. Hirsh, Receptor-mediated endocytosis in the Caenorhabditis elegans oocyte. Mol Biol Cell, 1999. 10(12): p. 4311–26.

53. Seah, N.E., et al., Autophagy-mediated longevity is modulated by lipoprotein biogenesis. Autophagy, 2016. 12(2): p. 261–72.

54. Steinbaugh, M.J., et al., Lipid-mediated regulation of SKN-1/Nrf in response to germ cell absence. Elife, 2015. 4.

55. Koyuncu, S., et al., Rewiring of the ubiquitinated proteome determines ageing in C. elegans. Nature, 2021. 596(7871): p. 285–290.

56. Watts, J.S., et al., New Strains for Tissue-Specific RNAi Studies in Caenorhabditis elegans. G3 (Bethesda), 2020.

57. Camara, H., et al., Tissue-specific overexpression of systemic RNA interference components limits lifespan in C. elegans. Gene, 2024. 895: p. 148014.

58. Weisman, A.S., N.M. Fisher, and C.P. Hunter, Efficient iterative CRISPR/Cas9 editing using sid-1 co-conversion and feeding RNAi in Caenorhabditis elegans. G3 (Bethesda), 2025. 15(8).

59. Babin, P.J., et al., Apolipophorin II/I, apolipoprotein B, vitellogenin, and microsomal triglyceride transfer protein genes are derived from a common ancestor. J Mol Evol, 1999. 49(1): p. 150–60.

60. Lapierre, L.R., et al., Amino acid sequences within the beta1 domain of human apolipoprotein B can mediate rapid intracellular degradation. J Lipid Res, 2004. 45(2): p. 366–77.

61. Cecchettini, A., et al., Vitellin cleavage products are proteolytically degraded by ubiquitination in stick insect embryos. Micron, 2003. 34(1): p. 39–48.

62. Kumar, A.V., et al., Lipid droplets modulate proteostasis, SQST-1/SQSTM1 dynamics, and lifespan in C. elegans. iScience, 2023. 26(10): p. 107960.

63. Bohnert, K.A. and C. Kenyon, A lysosomal switch triggers proteostasis renewal in the immortal C. elegans germ lineage. Nature, 2017. 551(7682): p. 629–633.

64. Eroglu, M., et al., Noncanonical inheritance of phenotypic information by protein amyloids. Nat Cell Biol, 2024. 26(10): p. 1712–1724.

65. Kern, C.C., et al., C. elegans feed yolk to their young in a form of primitive lactation. Nat Commun, 2021. 12(1): p. 5801.

66. Altin, S., et al., Ubiquitin precursor with C-terminal extension promotes proteostasis and longevity. Mol Cell, 2025. 85(19): p. 3677–3693 e7.

67. Papaevgeniou, N. and N. Chondrogianni, The ubiquitin proteasome system in Caenorhabditis elegans and its regulation. Redox Biol, 2014. 2: p. 333–47.

68. Johnston, S.C., et al., Structural basis for the specificity of ubiquitin C-terminal hydrolases. EMBO J, 1999. 18(14): p. 3877–87.

69. Mattiroli, F. and L. Penengo, Histone Ubiquitination: An Integrative Signaling Platform in Genome Stability. Trends Genet, 2021. 37(6): p. 566–581.

70. Mark, K.G. and M. Rape, Ubiquitin-dependent regulation of transcription in development and disease. EMBO Rep, 2021. 22(4): p. e51078.

71. Rangaraju, S., et al., Suppression of transcriptional drift extends C. elegans lifespan by postponing the onset of mortality. Elife, 2015. 4: p. e08833.

72. Janes, J., et al., Chromatin accessibility dynamics across C. elegans development and ageing. Elife, 2018. 7.

73. Brenner, S., The genetics of Caenorhabditis elegans. Genetics, 1974. 77(1): p. 71–94.

74. Love, M.I., W. Huber, and S. Anders, Moderated estimation of fold change and dispersion for RNA-seq data with DESeq2. Genome Biol, 2014. 15(12): p. 550.

75. Holdorf, A.D., et al., WormCat: An Online Tool for Annotation and Visualization of Caenorhabditis elegans Genome-Scale Data. Genetics, 2020. 214(2): p. 279–294.

76. Gu, Z., R. Eils, and M. Schlesner, Complex heatmaps reveal patterns and correlations in multidimensional genomic data. Bioinformatics, 2016. 32(18): p. 2847–9.

