## Supplemental Figures and Tables for "Validation of tissue-specific RNAi systems in *C. elegans* reveals a converging role for polyubiquitin UBQ-1/UBC in vitellogenin metabolism and lifespan"

### Figure S1

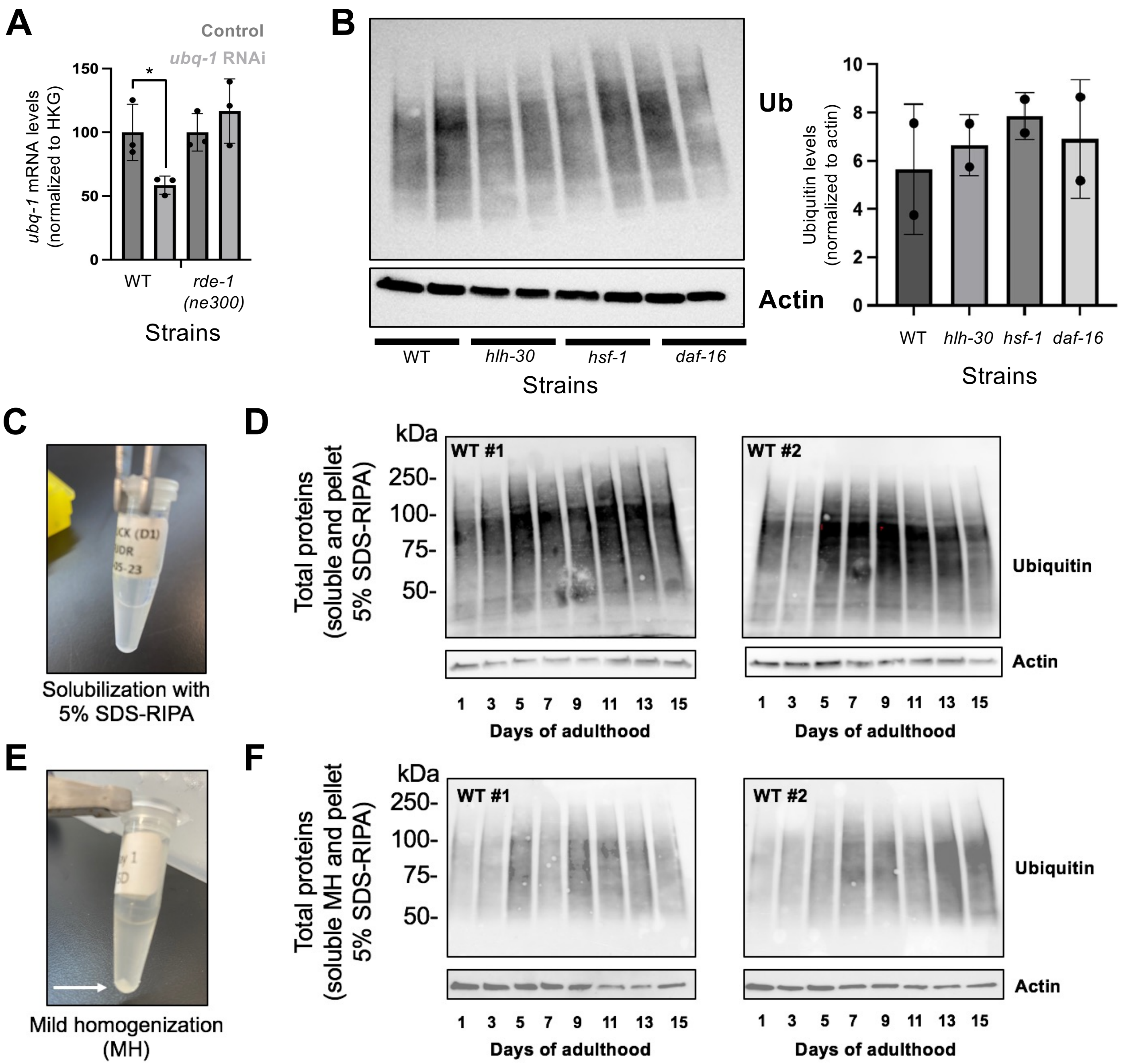


**Figure S1. Polyubiquitinated protein levels remain elevated throughout lifespan in *C. elegans***
**(A)** qPCR analysis of the expression of *ubq-1* in Day 4 wild type animals and *rde-1(ne300)* mutants fed control bacteria or bacteria expressing dsRNA against *ubq-1* from Day 1 to Day 4 of adulthood at 20°C, n=3, ±SD *t-test *:* P<0.05. Normalization was done with housekeeping (HKG) genes *act-1* and *cdc-42.* **(B)** Immunoblotting of polyubiquitinated proteins in Day 1 wild type animals, as well as *hlh-30(tm1978), hsf-1(sy441)* and *daf-16(mu16)* mutants, biological replicates ±SD. **(C)** Visualization of homogenate post-centrifugation after the first solubilization with 5% SDS-RIPA buffer (Day 1 sample). (**D)** Immunoblotting of total ubiquitinated proteins and actin in wild-type nematodes (#1, Kenyon lab N2CK, #2, Tuck lab N2ST) from Day 1 to Day 15 of adulthood, using 5% SDS-RIPA (both soluble and pellet fractions). **(E)** Visualization of homogenate post-centrifugation after the first solubilization with the mild homogenization (MH) buffer from Koyuncu et al. (Day 1 sample – white arrow highlights noticeable pellet). **(F)** Immunoblotting of total ubiquitinated proteins and actin in wild-type nematodes (#1, Kenyon lab, #2, Tuck lab) from Day 1 to Day 15 of adulthood using mild homogenization used in Koyuncu *et al.* (MH) for soluble fraction and 5% SDS-RIPA for pellet.

### Figure S2

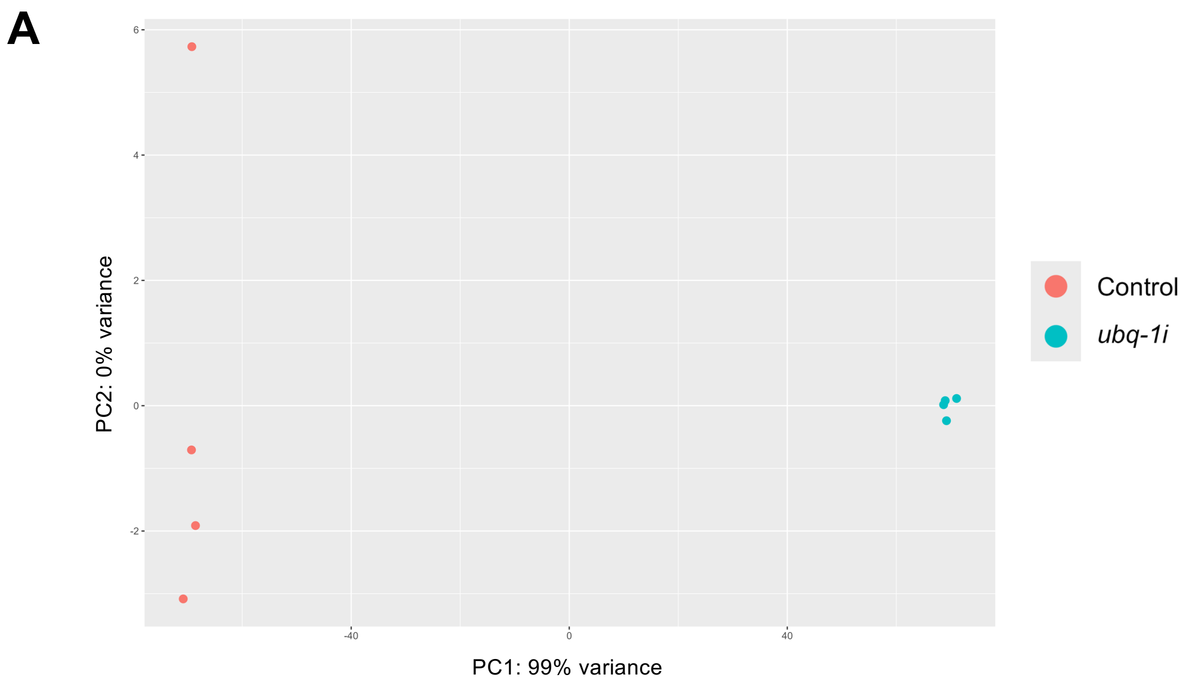


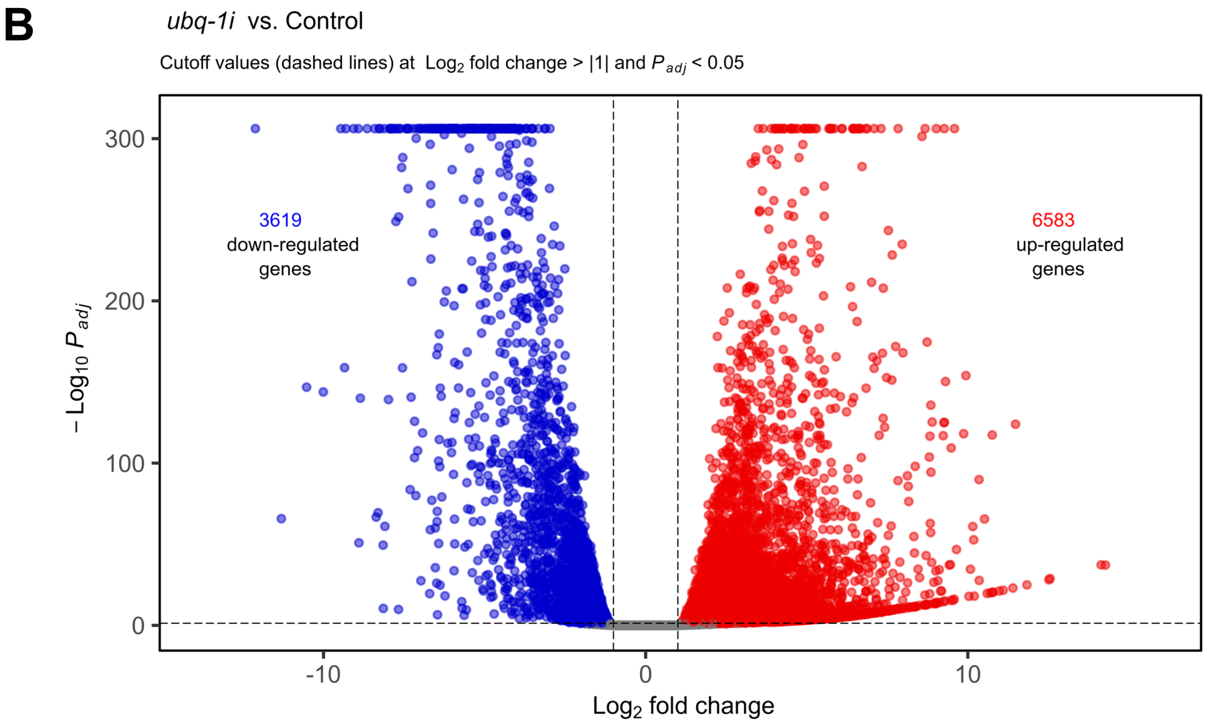


**Figure S2. Loss of *ubq-1* results in major transcriptional changes.**
**(A)** PCA analysis of samples used for RNA sequencing of mRNA from Day 4 wild type animals fed control bacteria or bacteria expressing *ubq-1* from Day 1 of adulthood at 20°C. **(B)** Volcano plot representation of differentially expressed genes between control and *ubq-1* silenced animals.

**Table S1**. Strains of *C. elegans* used in the study.

| Strain | Genotype | Mutation(s) | Lab |
|---|---|---|---|
| N2CK | Wildtype | - | Kenyon Lab |
| N2ST | Wildtype | - | Tuck Lab |
| CF512 | *fer-15(b26) II ;fem-1(hc17) IV* | Point mutation, Premature stop codon | Kenyon Lab |
| CF1038 | *daf-16(mu86) I* | Premature stop codon | Kenyon Lab |
| CF1903 | *glp-1(e2144) III* | L929F | Kenyon Lab |
| CF2495 | hsf-1(sy441) I | Point mutation | Kenyon Lab |
| DH1033 | *sqt-1(sc103) II*; bIs [vit-2::GFP + *rol-6 (su1006)*] *X* | Premature stop codon | Grant Lab |
| HC196 | *sid-1(qt9) V* | Premature stop codon (Q145*) | Hunter Lab |
| LRL1 | *hlh-30 (tm1978) IV* | Deletion mutation | NBRP |
| MAH85 | *glp-1(e2144) III. sid-1(qt9) V* | L929F (*glp-1*), Q145* (*sid-1*) | Hansen Lab |
| NL3321 | *sid-1(pk3321) V* | D130N | Plasterk Lab |
| WM27 | *rde-1(ne219) V* | E414K | Mello Lab |
| WM45 | *rde-1(ne300) V* | Premature stop codon (Q824*) | Mello Lab |
| **Strain** | **Genotype** | **Targeted tissue (Other tissues)** | **Lab** |
| AGD635 | *sid-1(qt9) V*; uthEx236[(*gly-19p*::tomato::54UTR), (*gly-19p*::sid-1(+)::unc-54UTR)] | Intestine (Hypodermis) | Dillin Lab |
| AGD637 | *sid-1(qt9) V*; uthIn201[(*rab-3p*::tomato::unc-54UTR), (*rab-3p*::sid-1(+)::unc-54UTR)] | Neurons | Dillin Lab |
| AGD639 | *sid-1(qt9) V*; uthEx238[(*myo-3p*::tomato::54UTR), (*myo-3p*::sid-1(+) unc-54 UTR)] | Muscle | Dillin Lab |
| AMJ345 | *rde-1(ne219) V*; jamSi2 [mex-5p::rde-1(+)] II. mex-5p::rde-1(+) inserted into ttTi5605 on LG II. | Germline + Intestine | Jose Lab |
| DCL569 | *sid-1*(mkc36) *V*; Si13 [sun-1p::rde-1::sun-1 3'UTR + unc-119(+)] II | Germline | Chen Lab |
| IG1839 | *rde-1(ne300) V*; frSi17 [mtl- 2p::rde-1 3'UTR] II. frIs7 [nlp-29p::GFP + col-12p::DsRed] IV. | Intestine | Watts Lab |
| IG1849 | *rde-1(ne300) V*; frSi21 [col-62p::rde-1 3'UTR] II. frIs7 [nlp-29p::GFP + col-12p::DsRed] IV | Hypodermis | Watts Lab |
| MAH677 | *sid-1(qt9) V*; sqIs71 [rgef-1p::GFP + rgef-1p::sid-1] | Neurons | Hansen Lab |
| NR222 | *rde-1(ne219) V*; kzIs9 [(pKK1260) lin-26p::NLS::GFP + (pKK1253) lin-26p::rde-1 + rol-6(su1006)] | Hypodermis (Int. Germ.) | Kalbuchi Lab |
| NR350 | *rde-1(ne219) V*; kzIs20 [hlh-1p::rde-1 + sur-5p::NLS::GFP] | Muscle | Kalbuchi Lab |
| VP303 | *rde-1(ne219) V*; kbIs7 [nhx-2p::rde-1 + rol-6(su1006)] | Intestine (Hypodermis) | Strange Lab |
| WM118 | *rde-1(ne300) V*; neIs9 [myo-3::HA::rde-1 + rol-6(su1006)] X | Muscle | Mello Lab |

### **Table S2.** Details of lifespan analyses performed at 20ºC

| Strain | Condition | Events observed/N | Mean Lifespan | Maximum Lifespan | % Difference | P value | Wild type reference |
|---|---|---|---|---|---|---|---|
| Wild type | control RNAi | 78/100 | 18.92 | 29 | | | - |
| | *ubq-1* | 97/100 | 5.86 | 7 | -69.03 | <0.001 | - |
| | control RNAi | 53/100 | 17.99 | 33 | | | - |
| | *ubq-1* | 93/100 | 6.70 | 10 | -62.76 | <0.001 | - |
| | control RNAi | 82/100 | 11.87 | 22 | | | - |
| | *ubq-1* | 91/100 | 6.37 | 8 | -46.33 | <0.001 | - |
| | control RNAi | 87/100 | 14.97 | 27 | | | - |
| | *ubq-1* | 95/100 | 6.46 | 8 | -56.85 | <0.001 | - |
| | control RNAi | 84/100 | 14.75 | 26 | | | - |
| | *ubq-1* | 91/100 | 6.37 | 8 | -56.81 | <0.001 | - |
| | control RNAi | 74/100 | 16.43 | 30 | | | - |
| | *ubq-1* | 96/100 | 6.49 | 8 | -60.50 | <0.001 | - |
| | control RNAi | 75/100 | 17.07 | 29 | | | - |
| | *ubq-1* | 94/100 | 5.81 | 8 | -65.96 | <0.001 | - |
| | control RNAi | 79/100 | 17.95 | 34 | | | - |
| | *ubq-1* | 97/100 | 5.84 | 8 | -67.46 | <0.001 | - |
| *rde-1 (ne300)* | control RNAi | 67/100 | 15.02 | 22 | | | |
| | *ubq-1* | 60/100 | 14.39 | 19 | -4.19 | 0.2488 | -62.76 |
| | control RNAi | 50/100 | 19.26 | 26 | | | |
| | *ubq-1* | 64/100 | 18.75 | 22 | -2.65 | 0.4073 | -69.03 |
| | control RNAi | 84/100 | 13.37 | 22 | | | |
| | *ubq-1* | 79/100 | 13.68 | 22 | 2.32 | 0.688 | -60.50 |
| *rde-1 (ne219)* | control RNAi | 75/100 | 19.54 | 29 | | | |
| | *ubq-1* | 72/100 | 11.48 | 19 | -41.25 | <0.001 | -69.03 |
| | control RNAi | 70/100 | 17.79 | 31 | | | |
| | *ubq-1* | 67/100 | 12.65 | 19 | -28.89 | <0.001 | -62.76 |
| | control RNAi | 79/100 | 19.83 | 31 | | | |
| | *ubq-1* | 79/100 | 10.29 | 19 | -48.11 | <0.001 | -46.33 |
| | control RNAi | 87/100 | 17.25 | 30 | | | |
| | *ubq-1* | 86/100 | 11.86 | 19 | -31.25 | <0.001 | -60.50 |
| *sid-1 (qt9)* | control RNAi | 76/100 | 16.46 | 29 | | | |
| | *ubq-1* | 83/100 | 10.89 | 17 | -33.84 | <0.001 | -60.50 |
| | control RNAi | 79/100 | 17.88 | 29 | | | |
| | *ubq-1* | 93/100 | 10.15 | 17 | -43.23 | <0.001 | -65.96 |
| | control RNAi | 86/100 | 18.62 | 31 | | | |
| | *ubq-1* | 90/100 | 10.00 | 17 | -46.30 | <0.001 | -67.46 |
| *sid-1 (pk3321)* | control RNAi | 86/100 | 17.19 | 24 | | | |
| | *ubq-1* | 43/100 | 8.69 | 11 | -49.45 | <0.001 | -69.03 |
| | control RNAi | 72/100 | 13.99 | 22 | | | |
| | *ubq-1* | 42/100 | 9.67 | 12 | -30.88 | <0.001 | -62.76 |
| | control RNAi | 67/100 | 13.12 | 22 | | | |
| | *ubq-1* | 83/100 | 7.68 | 12 | -41.46 | <0.001 | -60.50 |
| *glp-1 (e2144); sid-1 (qt9)* | control RNAi | 75/100 | 18.42 | 29 | | | |
| | *ubq-1* | 80/100 | 12.70 | 19 | -31.05 | <0.001 | -69.03 |
| | control RNAi | 79/100 | 15.09 | 24 | | | |
| | *ubq-1* | 72/100 | 11.68 | 15 | -22.60 | <0.001 | -62.76 |
| | control RNAi | 87/100 | 15.68 | 29 | | | |
| | *ubq-1* | 95/100 | 11.75 | 22 | -25.06 | <0.001 | -56.81 |

**Table S3.** Details of lifespan analyses performed at 25ºC

| Strain | Condition | Events observed/N | Mean Lifespan | Maximum Lifespan | % Difference | P value | Wild type reference |
|---|---|---|---|---|---|---|---|
| Wild type | control RNAi | 50/100 | 12.78 | 21 | | | - |
| | *ubq-1* | 92/100 | 6.57 | 7 | -48.60 | <0.001 | - |
| | control RNAi | 76/100 | 12.60 | 21 | | | - |
| | *ubq-1* | 97/100 | 4.92 | 8 | -60.95 | <0.001 | - |
| | control RNAi | 90/100 | 13.06 | 22 | | | - |
| | *ubq-1* | 97/100 | 5.06 | 8 | -61.26 | <0.001 | - |
| | control RNAi | 74/100 | 14.09 | 25 | | | - |
| | *ubq-1* | 97/100 | 6.56 | 8 | -52.90 | <0.001 | - |
| | control RNAi | 73/100 | 12.04 | 25 | | | - |
| | *ubq-1* | 88/100 | 6.30 | 7 | -47.67 | <0.001 | - |
| | control RNAi | 72/100 | 11.7 | 24 | | | - |
| | *ubq-1* | 96/100 | 5.05 | 7 | -56.84 | <0.001 | - |
| *rde-1 (ne300)* | control RNAi | 89/100 | 9.46 | 15 | | | |
| | *ubq-1* | 83/100 | 11.27 | 17 | 19.13 | <0.001 | -52.90 |
| | control RNAi | 74/100 | 10.56 | 17 | | | |
| | *ubq-1* | 66/100 | 12.05 | 21 | 14.11 | 0.0022 | -48.60 |
| | control RNAi | 78/100 | 11.36 | 21 | | | |
| | *ubq-1* | 84/100 | 11.85 | 23 | 4.31 | 0.4435 | -47.67 |
| *rde-1 (ne219)* | control RNAi | 82/100 | 12.9 | 26 | | | |
| | *ubq-1* | 86/100 | 13.2 | 26 | 2.32 | 0.8953 | -52.90 |
| | control RNAi | 84/100 | 13.05 | 21 | | | |
| | *ubq-1* | 79/100 | 13.69 | 24 | 4.90 | 0.3982 | -48.60 |
| | control RNAi | 88/100 | 12.82 | 22 | | | |
| | *ubq-1* | 90/100 | 12.31 | 22 | -3.98 | 0.616 | -60.95 |
| *sid-1 (qt9)* | control RNAi | 84/100 | 12.89 | 24 | | | |
| | *ubq-1* | 85/100 | 9.34 | 17 | -27.54 | <0.001 | -48.60 |
| | control RNAi | 71/100 | 13.87 | 25 | | | |
| | *ubq-1* | 69/100 | 10.73 | 18 | -22.64 | <0.001 | -52.90 |
| | control RNAi | 90/100 | 11.25 | 22 | | | |
| | *ubq-1* | 88/100 | 9.39 | 17 | -16.53 | <0.001 | -56.84 |
| *sid-1 (pk3321)* | control RNAi | 95/100 | 10.00 | 20 | | | |
| | *ubq-1* | 68/100 | 6.28 | 8 | -37.20 | <0.001 | -60.95 |
| | control RNAi | 67/100 | 9.45 | 20 | | | |
| | *ubq-1* | 58/100 | 6.18 | 9 | -34.60 | <0.001 | -47.67 |
| | control RNAi | 87/100 | 9.36 | 22 | | | |
| | *ubq-1* | 85/100 | 7.43 | 10 | -20.62 | <0.001 | -56.84 |
| *glp-1 (e2144); sid-1 (qt9)* | control RNAi | 93/100 | 11.85 | 26 | | | |
| | *ubq-1* | 94/100 | 9.85 | 17 | -16.88 | 0.0161 | -52.90 |
| | control RNAi | 86/100 | 11.27 | 21 | | | |
| | *ubq-1* | 92/100 | 10.81 | 21 | -4.08 | 0.3815 | -48.60 |
| | control RNAi | 97/100 | 10.33 | 20 | | | |
| | *ubq-1* | 87/100 | 9.74 | 16 | -5.71 | 0.1325 | -61.26 |
| | control RNAi | 99/100 | 14.28 | 25 | | | |
| | *ubq-1* | 100/100 | 10.76 | 16 | -24.65 | <0.001 | -56.84 |

**Table S4.** Primers used in the study.

| Primers used for qPCR | | | |
|---|---|---|---|
| **Primer** | **Direction** | **Sequence (5’ to 3’)** | **Target** |
| **LL1** | Forward | CTACGAACTTCCTG ACGGACAAG | *act-1* |
| **LL2** | Reverse | CCGGCGGACTCCAT ACC | *act-1* |
| **LL9** | Forward | CTGCTGGACAGGAAG ATTACG | *cdc-42* |
| **LL10** | Reverse | CTCGGACATTCTCGA ATGAAG | *cdc-42* |
| **LL667** | Forward | TCCGAGGAGGAATG CAATC | *ubq-1* |
| **LL668** | Reverse | TGGCCTTCACGTTC TCAATAG | *ubq-1* |
